# The allosteric mechanism leading to an open-groove lipid conductive state of the TMEM16F scramblase

**DOI:** 10.1101/2022.05.23.493170

**Authors:** George Khelashvili, Ekaterina Kots, Xiaolu Cheng, Michael V Levine, Harel Weinstein

**Author notes:** **Corresponding Author:** Address: Department of Physiology and Biophysics, Weill Cornell Medicine, 1300 York Avenue, room LC-501C, New York, NY, 10065.

## Abstract

TMEM16F is a Ca^2+^-activated phospholipid scramblase in the TMEM16 family of membrane proteins. Unlike other TMEM16s exhibiting a membrane-exposed hydrophilic groove that serves as a translocation pathway for lipids, the experimentally determined structures of TMEM16F shows the groove in a closed conformation even under conditions of maximal scramblase activity. Here we describe the analysis of ∼400 µs all-atom ensemble molecular dynamics (MD) simulations of the TMEM16F dimer that revealed an allosteric mechanism leading to an open-groove state of the protein that is competent for lipid scrambling. The grove opens into a continuous hydrophilic conduit that is highly similar in structure to that seen in other activated scramblases. The allosteric pathway connects this opening to an observed destabilization of the Ca^2+^ ion bound at the distal site near the dimer interface, to the dynamics of specific protein regions that produces the open-groove state seen in our MD simulations to scramble phospholipids.

## INTRODUCTION

The TMEM16F belongs to the TMEM16 family of integral membrane proteins. The human genome encodes for 9 TMEM16 protein homologs that have been classified as Ca^2+^-activated phospholipid scramblases (PLS), or Cl^-^ channels (CaCC) ^1–7^. The scramblase members of the family can also function as non-selective ion channels. They regulate the exposure of phosphatidylserine (PS) lipids on the cell surface and play essential roles in fundamental physiological processes, from blood coagulation, bone formation and cell fusion to membrane repair and immune response^8^. Dysfunction of TMEM16 PLS proteins is associated with genetically inherited disorders of muscle ^9,10^, bone^11–13^, blood^14,15^ and brain^16–19^, and mutations in TMEM16 PLS have been associated with various disease conditions ^5,20–24^. Specifically, mutations in human TMEM16F (hTMEM16F) PLS are responsible for Scott syndrome ^25^, a bleeding disorder caused by impairment of Ca^2+^-dependent externalization of PS lipids in activated platelets.

Much of what is known about structure/function relationships in TMEM16 PLS is inferred from breakthrough studies on fungal nhTMEM16 and afTMEM16 homologs ^1,2,26–33^, but for mammalian TMEM16 PLS such structure-based functional insights are only now starting to emerge ^6,34,35^. These insights have led to seemingly contradictory inferences, and it is still unclear to what extent the molecular mechanisms underlying activity or regulation of TMEM16 PLS are similar between mammalian and fungal homologs, or even among different mammalian ones. Thus, the X-ray and cryo-electronmicroscopy (cryo-EM) structures of Ca^2+^-bound fungal nhTMEM16 and afTMEM16 proteins ^27,36^ have revealed a common homo-dimeric fold, in which the ten transmembrane helices (TMs) of each protomer generate a hydrophilic groove facing the membrane on the side opposite to the dimer interface (Figure 1). This membrane-facing groove is lined by TMs 3-7 and connects (through TMs 6-7) to a pair of bound Ca^2+^ ions. In the absence of Ca^2+^ ions, the groove is occluded from the membrane as TM4 and TM6 helices reposition to close the groove. Structural, functional and computational experiments on the fungal TMEM16 PLS ^3,37,31,32,28,29,33^ revealed that in the presence of Ca^2+^ ions, the groove region in these scramblases can sample a wide range of conformations, such as an “ion-conducting” intermediate state (non-permissive to lipids but wide enough to conduct small size ions); a “membrane-exposed” open state (with an overall open conformation of the groove but still sufficiently narrow at the extracellular (EC) side to restrict the passage of lipid headgroups due to a constriction formed by the polar interaction network between TMs 3, 4, and 6); and a “lipid-conductive” state (in which the constriction is dissolved and the groove is transformed into a continuous hydrophilic conduit that allows the passage of lipids). Overall, these insights support the ‘credit card’ model mechanism for lipid scrambling ^38^ by the TMEM16 PLS, whereby lipids traverse the bilayer by populating the hydrophilic groove pathway with their headgroups while keeping their hydrophobic tails perpendicular to the groove axis, in the bilayer environment.

**Figure 1:**
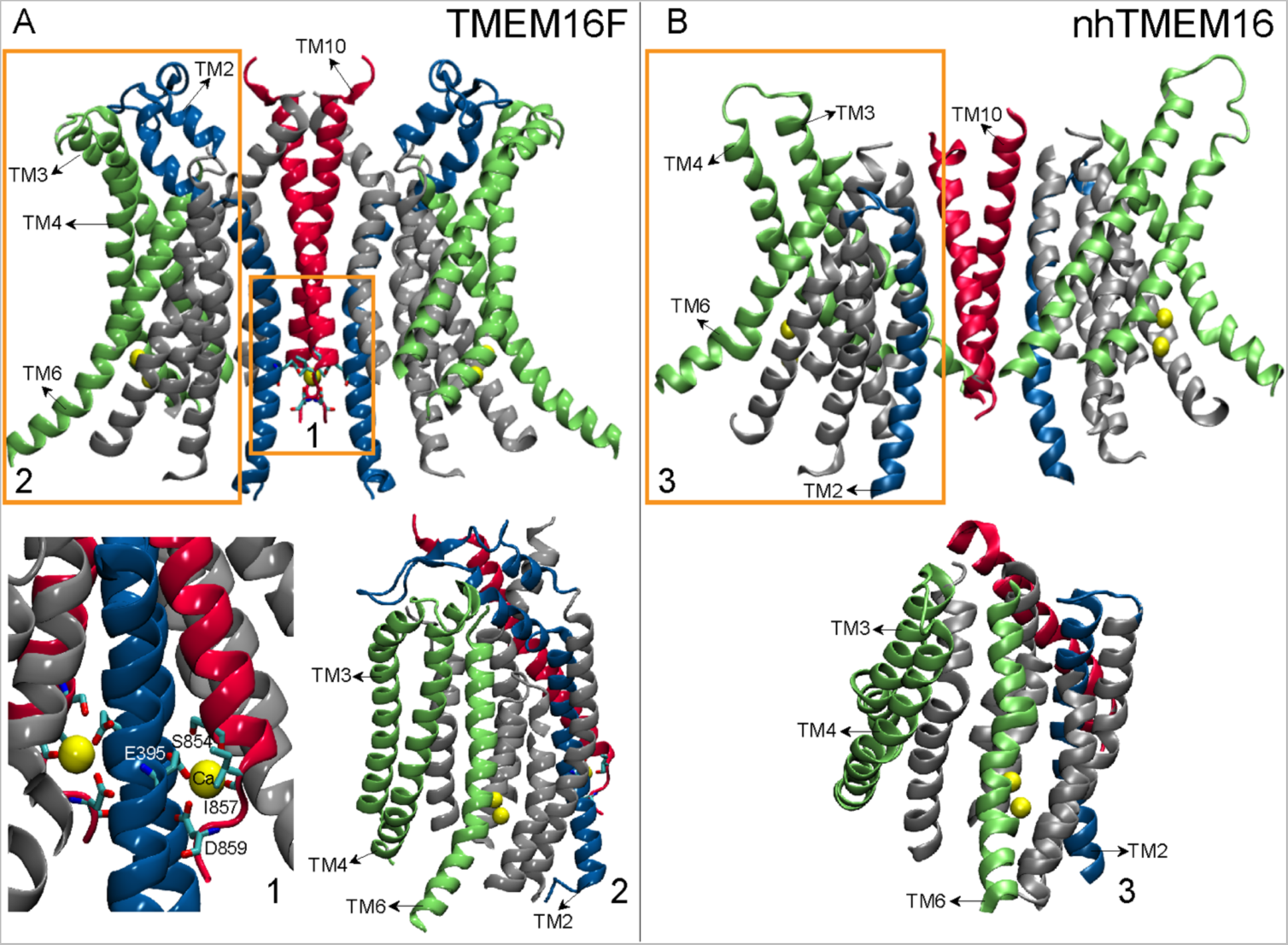
Distinguishing structural motifs of TMEM16F PLS are highlighted by a comparison of mTMEM16F (PDBID 6QP6), in Panel A (*left*) and nhTMEM16 (PDBID 6QM9) in Panel B (*right*). Most of the intracellular and extracellular loop regions have been removed for clarity. In Panel A, rectangle 1 highlights the third (distal) Ca^2+^ binding site at the dimer interface of mTMEM16F. Shown in licorice and labeled are the 4 residues coordinating the Ca^2+^ ion in its binding site (E395 on TM2, and S854, I857, and D859 on TM10). Rectangle 2 in Panel A and Rectangle 3 in Panel B highlight the groove regions in mTMEM16F and nhTMEM16, respectively. Shown are TMs 3, 4, and 6 (in green) lining the groove region, TM10 (in red) and TM2 (in blue). Note that the linker region connecting TMs 1 and 2 in the mTMEM16F model reaches towards the groove region and is positioned close to the extracellular end of TM4 helix (see also Figure S1).

More recent structure/function studies on mammalian hTMEM16K PLS ^35^ have provided further support for this mechanism, as the Ca^2+^-bound groove in hTMEM16K was found to assume conformations wide enough to constitute a lipid translocation pathway. Indeed, MD simulations suggested that such an open-groove state in hTMEM16K is necessary for scramblase activity ^35^. However, structural investigations of mouse TMEM16F (mTMEM16F) ^6,34^ showed in striking contrast to the fungal TMEM16 PLS and the hTMEM16K, the groove in mTMEM16F remains closed even under conditions of maximal activation (i.e., Ca^2+^-bound and in the presence of highly anionic phosphatidylinositol 4,5-bisphosphate (PIP_2_) lipids). This led to the proposal that mTMEM16F PLS may function according to a different paradigm ^34,39^, where lipid permeation occurs outside a closed scrambling pathway.

Comparison of existing structural data for the different TMEM16 PLS identifies two intriguing structural characteristics of mTMEM16F that together distinguish it from the fungal and hTMEM16K PLS (see Figure 1 and Figure S1): (1)-the closed groove in mTMEM16F on the EC side appears to be stabilized not only by TM4-TM6 interactions (referred here as TM4-TM6 interface as seen in the fungal and hTMEM16K PLS), but also by interactions between the EC end of TM4 helix and the extended helix-loop-helix motif on the EC end of TM2 (TM4-TM2 interface); and (2)-the mTMEM16F structure contains an additional, third Ca^2+^ ion bound at a “distal Ca^2+^ binding site” in the dimer interface region. The ion is stabilized at this site by the sidechains of two negatively charged residues – E395 on TM2, and D859 on TM10 – as well as by two backbone carbonyls from residues S854 and I857 in TM10. While the analogous Ca^2+^ ion binding site exists in the hTMEM16K structures as well, the unique structural feature of mTMEM16F is that the intracellular (IC) end of TM2 coordinates the third Ca^2+^ ion while the EC end of the same helix participates in the interactions with TM4 to provide stability to the closed groove structure. These characteristics present an intriguing possibility that the distal Ca^2+^ ion site in mTMEM16F may be allosterically coupled to the EC side of the groove. Indeed, Ca^2+^ binding to the analogous site in the TMEM16A CaCC site, has been shown to facilitate TMEM16A channel opening ^40^.

We therefore reasoned that, dynamics of the Ca^2+^ ion in the distal binding site may allosterically affect conformational dynamics of the groove region in mTMEM16F and lead to its opening for lipid scrambling. To test this hypothesis we carried out massive, ∼400 µs-long, all-atom ensemble molecular dynamics (MD) simulations of mTMEM16F. The analysis of the MD data revealed a gradual opening of the groove region and its transformation into a continuous lipid conduit. This conformational transition severs the interactions along the interfaces between TM4 and TM6, and between TM4 and TM2. Importantly, the analysis of MD trajectory data with Markov State Model (MSM) and Transition Path Theory (TPT) approaches revealed that the unraveling of the TM4-TM2 interface was correlated in time with the destabilization of the Ca^2+^ ion in the distal binding site. To discern the mode of allosteric coupling between the ion binding site and the groove region we used N-body Information Theory (NbIT) analysis ^41^ and quantified it with the Thermodynamic Coupling Function (TCF) approach ^42,43^. These revealed the allosteric path of communication between the two regions and showed that the mTMEM16F groove conformation that permits lipid scrambling is indeed energetically stabilized by the allosteric communication. These results provide specific mechanistic details in a structural context for the allosteric mechanism leading to an open, lipid conductive state of the groove in mTMEM16F.

## METHODS

### Molecular constructs for molecular dynamics (MD) simulations

The all-atom MD simulations described in this work are based on the cryo-EM structure of the mTMEM16F protein (PDBID: 6QP6 ^6^). The missing segments from the structure (1–42, 150–186, 489–502, and 876–911) were modeled using Rosetta’s AbinitoRelax application ^44^. For the 876–911 segment, 1000 structures were generated and clustered. The most populated cluster (with clustering threshold of 4 Å) consisted of 315 structures, in all of which the 891–904 fragment was alpha helical. Based on the positioning of the analogous helical fragment in the structure of fungal homolog nhTMEM16 (PDBID 6QM5 ^45^), we positioned the 891–904 segment of mTMEM16F to interact with the N-terminal part of the opposite monomer (near residues 30-39, see Figure S2) in the starting structure of the simulations. The 428–444 missing stretch in the 6QP6 structure was modeled as a hairpin connecting TM2 and TM3 using Modeller v9 ^46^ with nhTMEM16 (6QM5) ^45^ and hTMEM16K(5OC9) ^35^ as structural templates. In the completed structure, the first 20 N-terminal 20 were excluded due to their high flexibility, so that the final model contained residues 21-911 (Figure S2). Protonation states of the titratable residues were predicted with Propka 3.1 ^47^, resulting in histidine 275, 655, and 818 modeled as HSE type, while the other histidine residues were modeled as HSD type.

Using the CHARMM-GUI web interface ^48^, the mTMEM16F model was embedded into a lipid membrane consisting of a 7:3 mixture of POPE (1-palmitoyl-2-oleoyl-sn-glycero-3-phosphoethanolamine) and POPG (1-palmitoyl-2-oleoyl-sn-glycero-3-phospho-(1’-rac-glycerol)) lipids, the same lipid composition as used in our previous simulations of nhTMEM16 PLS ^26,28,29^. The protein to lipid ratio was 1:1050. After adding a solvation box containing 150 mM K^+^Cl^-^ the total system contained ∼ 561,219 atoms.

### Atomistic MD simulations

The assembled system was subjected to a short equilibration run with NAMD 2.13^49^ using a standard set of equilibration scripts provided by CHARMM-GUI. After this initial equilibration, the velocities of all the atoms were randomly regenerated and the system was subjected to an extensive 3-stage adaptive ensemble MD simulation protocol (Figure S3). In Stage 1, the system was simulated in 300 independent replicates, each ∼200ns long. The analysis of the trajectories to assess the extent of conformational sampling (see Results) identified 48 frames for the next round of simulations (Figure S4A). In Stage 2, these 48 structures were run in 3 independent replicates (144 replicates in total), each for ∼1.5 µs. Another round of analysis further identified 8 trajectory frames from Stage 2 simulations to be considered for the next iteration (Figure S4B). Thus, in Stage 3 of ensemble simulations, these 8 frames were run in 18 independent replicates (144 replicates in total), each for ∼800ns. Overall, this multi-stage adaptive protocol resulted in a net sampling time of ∼400 µs.

The ensemble MD simulations were carried out with OpenMM 7.4 ^50^ and implemented PME for electrostatic interactions. The runs were performed at 310 K temperature, under NPT ensemble using semi-isotropic pressure coupling, and with 4fs integration time-step (with mass repartitioning). Monte Carlo barostat and Langevin thermostat were used to maintain constant pressure and temperature, respectively. Additional parameters for these runs included: “friction” set to 1.0/picosecond, “EwaldErrorTolerance” 0.0005, “rigidwater” True, and “ConstraintTolerance” 0.000001. For the van der Waals interactions we applied a cutoff distance of 12 Å, switching the potential from 10 Å.

For all simulations we used the latest CHARMM36m force-field for proteins and lipids ^51^, as well as the recently revised CHARMM36 force-field for ions which includes non-bonded fix (NBFIX) parameters ^52^.

### Dimensionality reduction with the tICA approach

To facilitate analysis of conformational dynamics in the simulations, we performed dimensionality reduction using tICA (time-lagged independent component analysis)^53^ as previously described ^28,29,54–57^. Briefly, in the tICA approach the MD simulation trajectories are used to construct two covariance matrices: a time-lagged covariance matrix (TLCM) *CTL(τ)=<X(t)XT(t+τ)>*, and the usual covariance matrix *C=<X(t)XT(t)>*, where *X(t)* is the data vector at time *t*, *τ* is the lag-time of the TLCM, and the symbol *<…>* denotes the time average. To identify the slowest reaction coordinates of the system, the following generalized eigenvalue problem is solved: *CTLV = CVΛ*, where *Λ* and *V* are the eigenvalue and eigenvector matrices, respectively. The eigenvectors corresponding to the largest eigenvalues define the slowest reaction coordinates. These reaction coordinates depend on the choice of data vector *X*, i.e., the choice of collective variables (CV). Here, to define the tICA space, we considered CVs that quantify conformational dynamics of the TM4-TM6 and TM4-TM2 interfaces of the mTMEM16F groove, as well as the coordination of the Ca^2+^ ion bound at the third binding site, referred throughout simply as the third Ca^2+^ ion (see Results, Figure 1). Thus, the following 3 sets of dynamic measures were extracted from the analysis of the MD trajectories to serve as CVs: 1) the C_α_-C_α_ distance between residue pairs 502-364, 510-604, 510-607, 514-608, 515-611, 517-608, 518-611, 518-612, 521-612, 522-619, and 525-616 (TM4-TM6 interface); 2) the C_α_-C_α_ distance between the residue pairs 497-360, and 499-363 (TM4-TM2 interface); 3) any minimum distance between the third Ca^2+^ ion and residues E395, S854, I857, and D859.

### Markov State Model (MSM) Construction

To perform MSM analysis, we used Python 2.7.14 scripts included in MSMBuilder software ^58,59^. Briefly, the 2D tICA space was discretized into 100 microstates using automated clustering *k-means* algorithm, and a transition probability matrix (TPM) was built ^58^. To ensure Markovian behavior, multiple TPMs were constructed for different time intervals between transitions (MSM lag times), and the relaxation timescales of the system were calculated as:

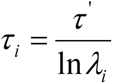

where *τ^’^* is the lag-time used for building the TPM, *λ_i_* denotes the *i*^th^ eigenvalue of the TPM, and *τ_i_* represents relaxation timescale (implied timescale) corresponding to the *i*^th^ relaxation mode of the system. The Markovian property of the TPM was established by verifying the independence of *τ_i_* from *τ^’^*. This analysis identified 160 ns as the lag-time at which the implied timescales begin to converge. Thus, the final MSM was built using *τ^’^* = 160 ns.

### Transition Path Theory (TPT) analysis

In order to identify the most probable pathways, we applied TPT analysis as previously described ^56^. Briefly, using a Robust Perron Cluster Analysis (PCCA+) algorithm ^60^, the 100 microstates within the tICA space were clustered into 16 macrostates based on their kinetic similarity. Using the Dijkstra graph theory algorithm ^61^ implemented in the MSMbuilder software, a flux matrix ^62^ was then constructed for macrostates, and the most probable pathways were identified as those with the highest flux between the starting macrostate and a final microstate.

### Thermodynamic Coupling Function (TCF) analysis

The TCF analysis was carried out as previously described ^42,43^. Briefly, given two CVs, 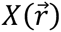 and 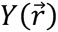, which are functions of only the atomic coordinates, 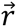, the TCF identified as ΔΔ*A*(*x*, *y*), is a function of specific values (*x*, *y*) of those CVs:

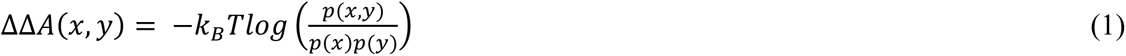

In the above, *k_B_* is the Boltzmann constant, T is the temperature, and *p(x,y), p(x)* and *p(y)* are, respectively, the observed joint and marginal probability distributions of molecular conformations. Here, 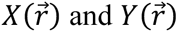 were calculated from the normalization of the first two tIC vectors as ^42^:

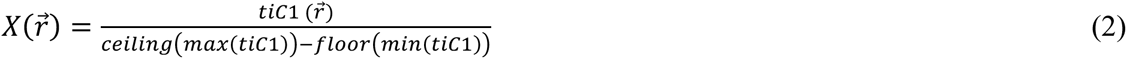

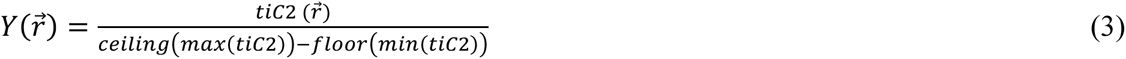

Each configuration in microstate *m* was given a weight:

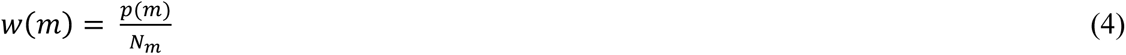

where *p*(*m*) represents the equilibrium weight of the microstate taken from the MSM, and *N_m_* is the number of configurations associated with this microstate. The 2D space of the first two tIC vectors was then divided into 100 equally spaced bins in each direction, and the joint probability density corresponding to a bin centred at (*x, y*) was estimated over the configuration probability density within that bin ^42^:

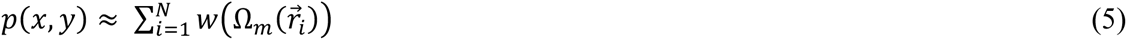

where 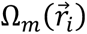 is the index of the microstate a given configuration resides in, and 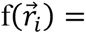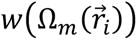 represents the probability density function over the configurations 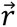. From Eq. (5) above, p(x) and p(y) were calculated by summing over the corresponding dimensions.

Estimation of confidence levels for each bin was performed using the bootstrapping procedure described previously ^42^.

### N-body Information Theory (NbIT) analysis

To quantify the allosteric coupling between the distal Ca^2+^-binding site and the mTMEM16F groove region we have applied N-body Information Theory (NbIT) analysis ^41^ to the trajectories of metastable MSM microstates which represent local energy minimum conformations on the tICA space (see Results). The NbIT method relies on the Information Theory representation of configurational entropy *H* calculated from the covariance matrix (***C***) of all atomic positions (**X**) in the protein:

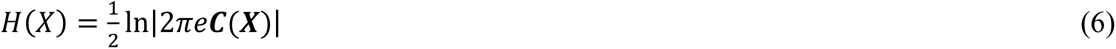

Total correlation (*TC*), defined as the total amount of information that is shared among *N* atoms in a set, was then quantified as:

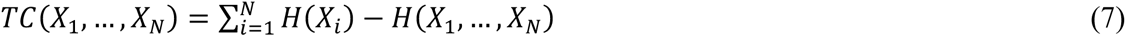

where *X_i_*-s represent components of 3N-dimensional vector corresponding to *x*, *y*, *z* atomistic coordinates of the atoms in the set, and *H*(*X*_1_, …, *X_N_*) is the joint entropy of the set.

Using *TC*, we then obtained coordination information (*CI*) which quantifies the amount of information shared by a set of variables of arbitrary size that is also shared with another variable:

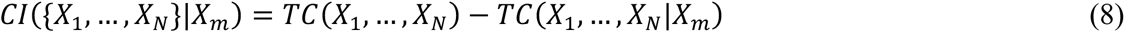

In the above, *TC*(*X_1_*, …, *X_N_*|*X_m_*) represents conditional total correlation between {*X_1_*, …, *X_N_*}, conditioning on *X_m_*.

From *CI*, we calculated the mutual coordination information (MCI), defined as the amount of coordination information that is shared between two residues and the same set:

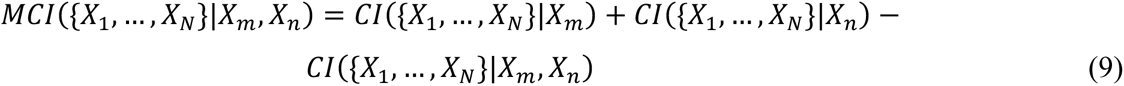

Here we have applied NbIT to all the non-hydrogen atoms of the mTMEM16F protein. First, we evaluated the *CI* values in various MSM metastable states between the distal Ca^2+^-binding site (S1 site: residues E395, S854, I857 and D859) as the *transmitter*, and the groove of mTMEM16F the *receiver* (see ^41^ for details). Several additional sites along the TM4-TM6 and the TM4-TM2 interfaces were also defined as *receivers*: S2 site: W359/Y502, S3 site: Q506/Q351; S4 site: Q351/Y502; S5 site: Y502/Q506; S6 site: M522/W619; S7 site: F518/K616; S8 site: S514/ Q608; S9 site: S514/F518/M522; S10 site: Q608/K616/W619. The *CI* results were further clustered with Fisher-Jenks algorithm (https://github.com/mthh/jenkspy) by the level of coordination into three groups: low (*CI* < 24%), average (24% ≤ *CI* ≤ 50%), and high (*CI* > 50%). The allosteric channels connecting the distal Ca^2+^-binding site with the TM4-TM2 and TM4-TM6 interfaces was identified by evaluating the *MCI* between the S1 site and the combined sites S2+S3+S4+S5 and S6+S7+S8 sites.

## RESULTS

### Opening of the mTMEM16F groove region is related to the restructuring of the distal Ca^2+^ binding site

A unique structural feature of mTMEM16F PLS, not observed in any available structural models of other TMEM16 scramblases, is the involvement of TM2 helix not only in coordination of the Ca^2+^ ion in its binding site at the dimer interface (distal Ca^2+^ binding site), but also in the apparent stabilization of the closed groove structure of mTMEM16F by interactions with TM4 (Figure 1). Based on the span of this structural feature we hypothesized that it may support an allosteric connection between the intracellularly located, distal Ca^2+^ binding site, in mTMEM16F and the extracellular end of the groove. To test this hypothesis, we set out to probe whether the dynamics of the distal Ca^2+^ binding site allosterically affects conformational changes in the groove region to yield its opening. As described in Methods, the massive MD simulations totaling ∼400 µs trajectory time were performed in a 3-stage adaptive protocol, in which each Stage is informed by the output from the previous one used to spawn swarms of multiple replicates (see Methods, Figure S3). In the course of these simulations we observed a gradual opening of the groove region of one of the protomers of mTMEM16F that allowed lipid scrambling, concomitant with a destabilization of the Ca^2+^ ion in the distal binding site (Supplemental Movies 1 and 2). To facilitate analysis of these conformational changes, we performed dimensionality reduction using tICA formulation as described in Methods. To this end, we extracted from the MD trajectories three sets of dynamic CVs to quantify (i)-conformational changes at the TM4-TM6 and TM4-TM2 interfaces of the groove, and (ii)-changes in the coordination of the Ca^2+^ ion in its binding site (Figure 2A, see Methods). The tICA transformation of the MD trajectory frames to the space of these variables showed that the first two tIC vectors (tIC1 and tIC2) represented >80% of the total dynamics in the system (Figure 2C). Evaluation of contributions of each of the CVs to these vectors revealed that tIC1 encoded mainly structural changes at the TM4-TM6 interface, whereas tIC2 mostly encoded structural changes at the TM4-TM2 interface and in the distal Ca^2+^ ion coordination. The projection of all the MD trajectory frames onto a 2D space of the first two tIC vectors is shown in Figure 3. Structural characteristics of the conformations sampled by the groove region and by the distal Ca^2+^ binding site in the simulations are indicated by molecular representations of the selected microstates shown in the snapshots provided in Figure 3. In Figure S5 we show the results of discretizing this space into microstates as described in Methods. Results of the quantification of various collective variables in these microstates are presented in the histograms in Figure S6.

**Figure 2:**
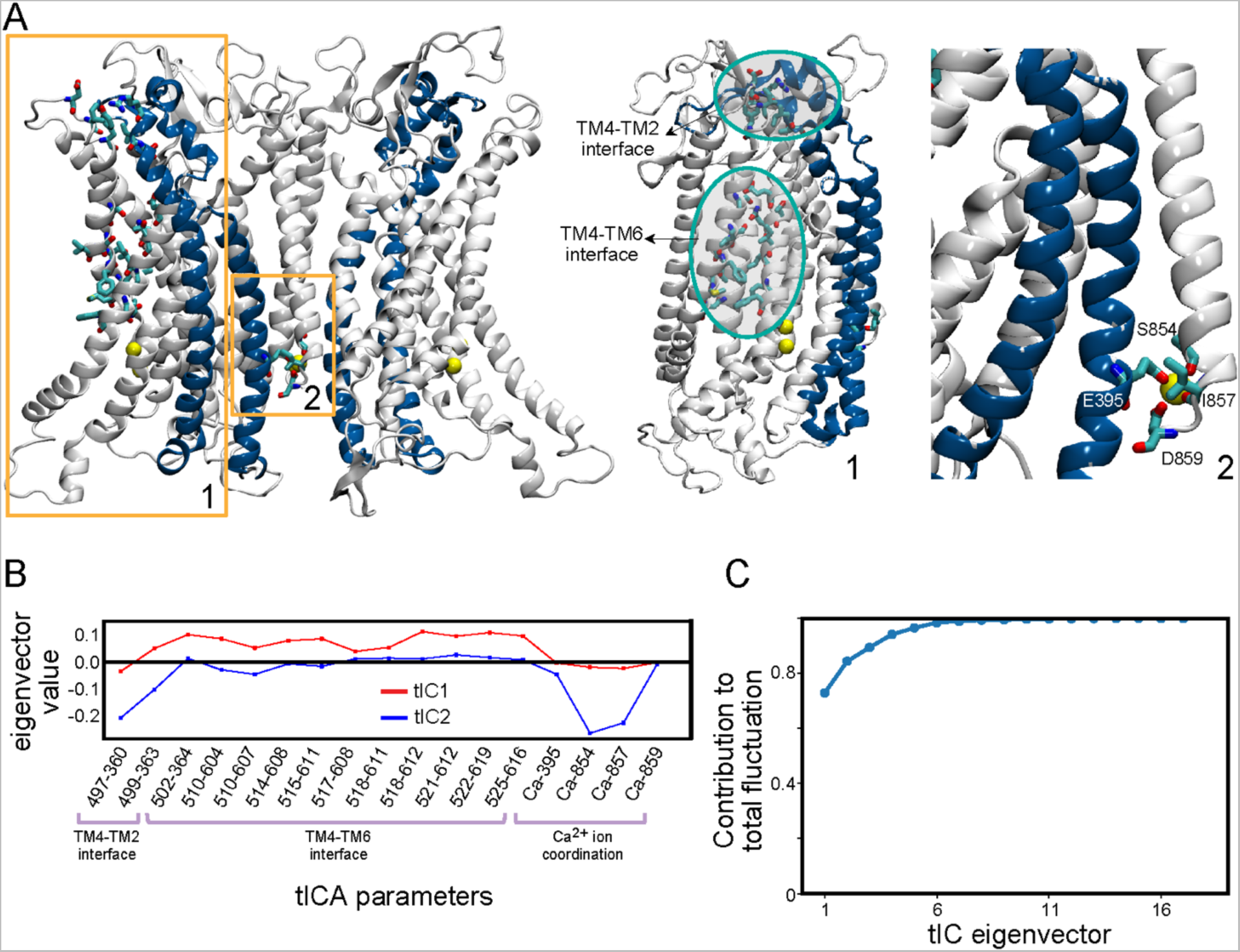
Description of the CVs used in the tICA dimensionality reduction analysis. (**A**) The three regions whose conformational changes are described with the chosen CVs are highlighted on the structure of the mTMEM16F model by the numbered rectangles: **1.** TM4-TM6 and TM4-TM2 interfaces of the groove (inset *i*); **2.** The distal Ca^2+^ binding site. In **1**, the specific residues whose pairwise distances throughout the trajectory served as CVs are shown rendered in licorice. In **2**, the four residues coordinating the Ca^2+^ ion (yellow sphere) in the binding site are shown rendered in licorice. (**B**) Contributions of the CVs used as tICA parameters to the tIC1 and tIC2 vectors. (C) Contribution of each tIC vector to the total representation of structural dynamics of the system.

**Figure 3:**
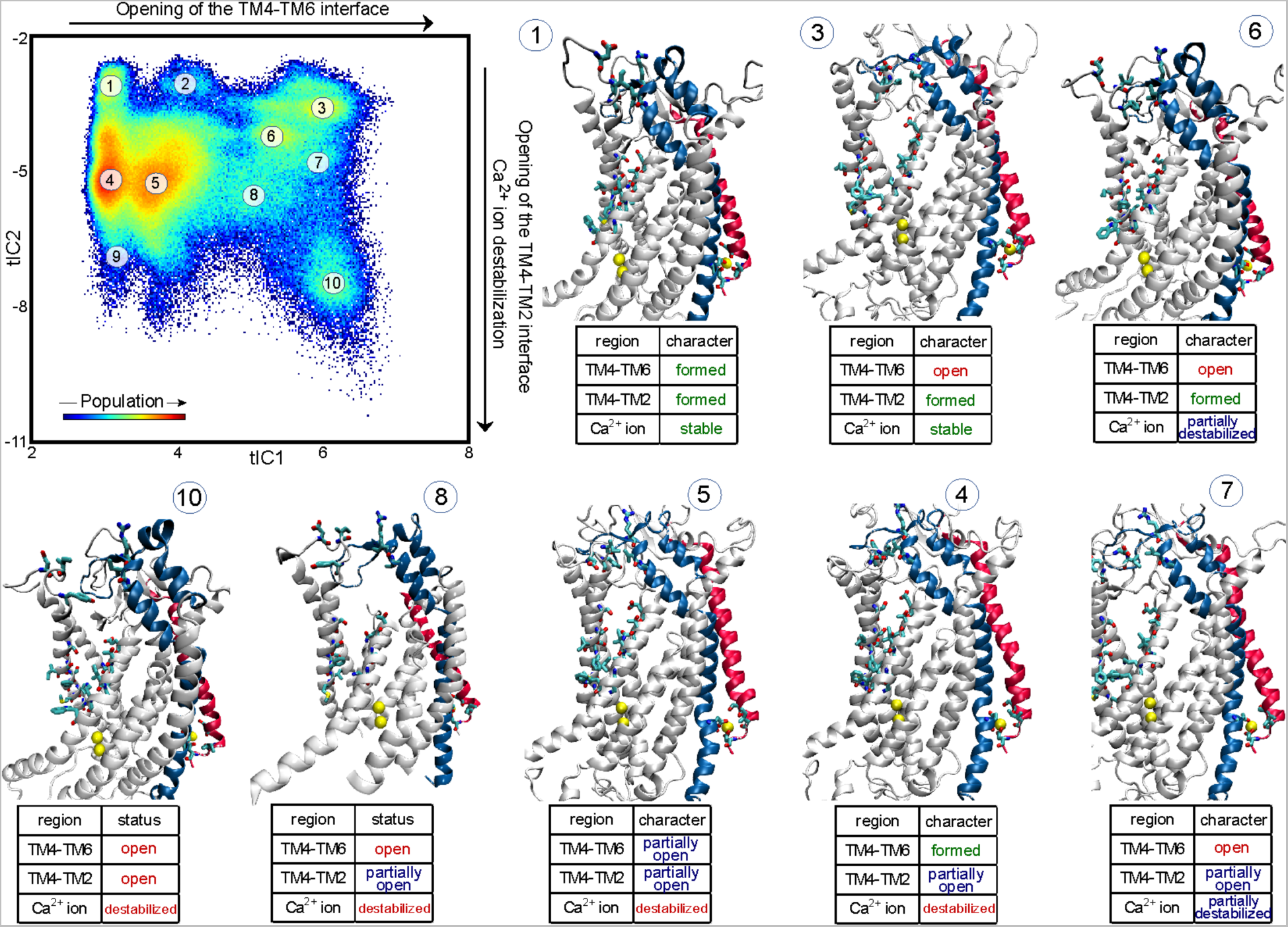
Structural interpretation of the tICA space features. Projection of all the MD trajectory frames from the adaptive ensemble simulations onto the 2D space spanned by the first two tIC vectors (see also Figure S3). The color map identifies the populations distribution of the different states of mTMEM16F in this tICA space. Representative structures from selected microstates indicated on the tICA map are shown in the molecule models. The table accompanying each model describes the characteristics of the critical regions (TM4-TM6, TM4-TM2, and the distal Ca^2+^ ion binding site) in the corresponding microstate. Only one protomer of mTMEM16F is shown. TMs 2 and 10 are colored in blue and red, respectively. The Ca^2+^ ions are shown as yellow spheres and the relevant residues are drawn in licorice as in Figure 2A. A more detailed structural quantification of the microstates is presented in Figure S6.

The extent of TM4-TM6 interface opening represented by moving along the first eigenvector, tIC1, in the 2D tICA space (Figures 3 and S6) can be divided into 3 parts: for small tIC1 values (microstates 1, 4, and 9), the TM4-TM6 interface of the groove is closed; for intermediate tIC1 values (microstates 2 and 5), it is partially open; and for large tIC1 values (microstates 3, 6, 7, 8, and 10), the TM4-TM6 interface is fully open. Movement along the second eigenvector, tIC2, in Figures 3 and S6 reflects the extent of opening of the TM4-TM2 interface. The 2D tICA space can again be considered as 3 regions: as tIC2 values decrease from -2 to -11, the system first visits conformations with closed TM4-TM2 interface (microstates 1, 2, 3, and 6), followed by conformations with partially open (microstates 4, 5, 7, and 8), and then fully open (microstates 9 and 10) TM4-TM2 interface. This fully open structure of the groove contains a continuous membrane-exposed pore whose dimensions are comparable to those measured in the open groove structure of nhTMEM16 ^36^ (see Figure S7).

A restructuring of the distal Ca^2+^ binding site is also occurring along tIC2. At values in the range from -2 to ∼ -4 (microstates 1-3 in Figures 3 and S6), the Ca^2+^ ion is stably bound in its site, coordinated by the sidechains of residues E395 and D859, as well as by backbone carbonyls of S854 and I857. But for tIC2 values < -4 (microstates 4-10), the Ca^2+^ ion is gradually destabilized as indicated by the loss of coordination with the backbone moieties of S854 and I857 (coordination with charged sidechains of E395 and D859 is maintained; see also Figure S8A-B). Concomitantly, the Ca^2+^ ion becomes more hydrated (see “Ca^2+^ hydration” histograms in Figure S6) and moves inward by ∼5Å while bringing with it the intracellular segment of TM2 (located below residue E395) and the terminal intracellular helical portion of TM10 (albeit to a lesser extent, see Figure S8C). These conformational changes on the intracellular side are accompanied by the rearrangements of the extracellular end of TM2 which moves away from TM4 as the Ca^2+^ ion becomes destabilized. Thus, the tICA analysis records the evolution of the system from the ensemble of cryo-EM like states (microstate 1 in Figure 3) – in which both TM4-TM6 and TM4-TM2 interfaces are closed and the distal Ca^2+^ binding site is intact – to the ensemble of conformations with both TM4-TM6 and TM4-TM2 interfaces of one of the monomers fully ruptured and the Ca^2+^ ion at the dimer interface destabilized and displaced from its binding site (microstate 10 in Figure 3).

To identify kinetic pathways leading to the groove opening and concomitant destabilization of the Ca^2+^ ion at the dimer interface, we built an MSM model on the microstates of the tICA space. As described in Methods, the microstates were lumped into 16 macrostates based on the MSM analysis (Figure S5). The TPT analysis performed on these macrostates then identified top state transition pathways connecting structural states in the tICA space (see Methods), and revealed a dominant pathway leading to the structural changes described above (Figure 4). Along this top pathway, the system first undergoes a transition from the ensemble of cryo-EM like states to the ensemble of conformations in which the TM4-TM6 interface of the groove is still intact, but the distal Ca^2+^ binding site is destabilized and the TM4-TM2 interface is partially open (in Figure 4, see transition from macrostate **a** to **b**). This is then followed by the transitions to the states in which the TM4-TM6 interface begins to gradually open while the distal Ca^2+^ binding site remains destabilized and TM4-TM2 interface – partially open (in Figure 4, see transitions from macrostate **b** to **c**, and **c** to **d**). Lastly, the system transitions to the ensemble of states in which both TM4-TM6 and TM4-TM2 interfaces of the groove are fully open and the distal Ca^2+^ binding site is destabilized (macrostate **d** to **e** transition in Figure 4). The other top pathways identified from the TPT analysis largely follow a similar sequence of kinetic steps in which Ca^2+^ destabilization precedes full opening of the groove.

**Figure 4:**
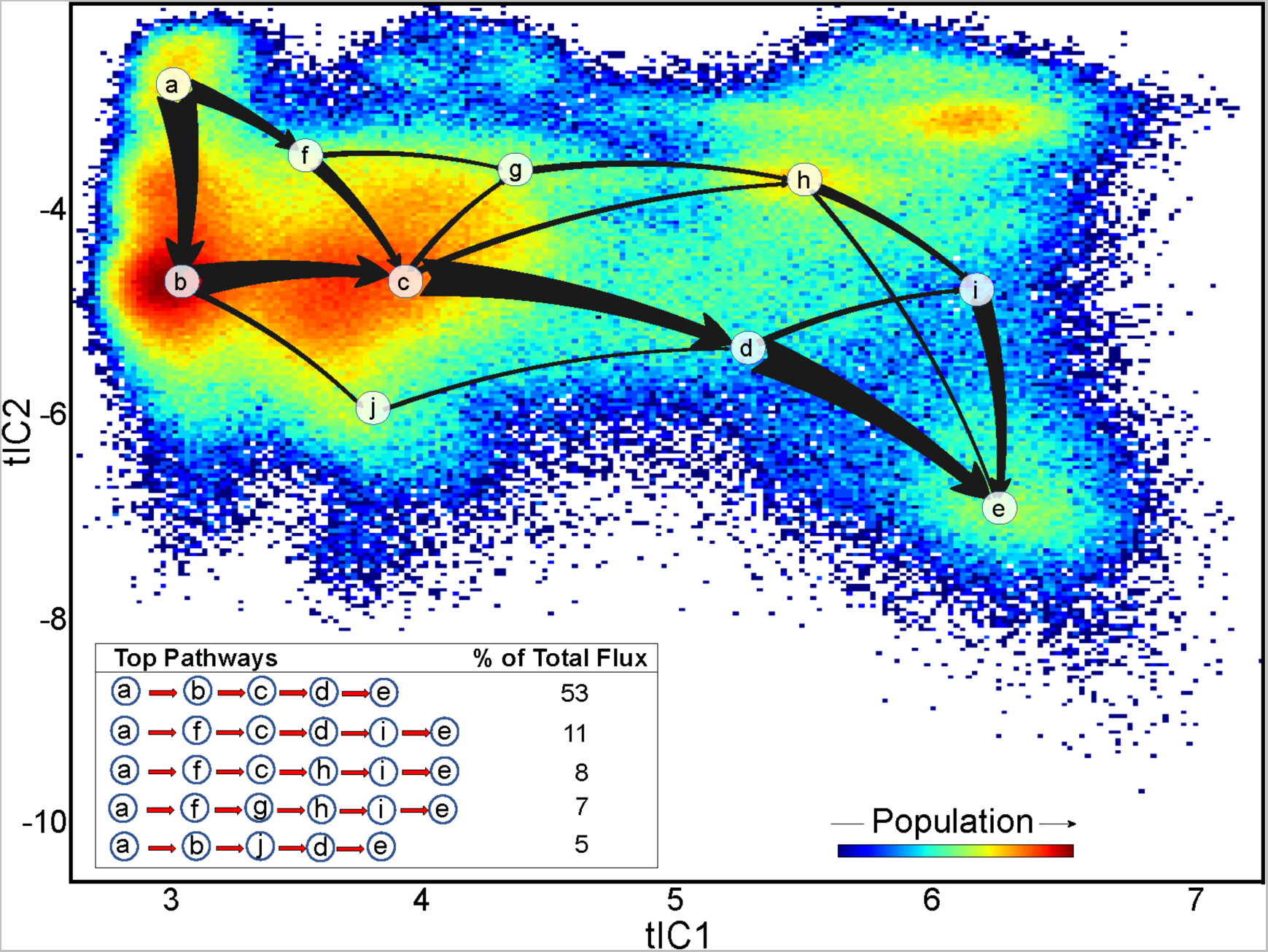
Kinetic pathways leading to groove opening states and destabilization of the distal Ca^2+^ ion at the dimer interface. The top 5 pathways identified from the TPT analysis to connect macrostate **a** (closed TM4-TM6 interface, closed TM4-TM2 interface, stably bound Ca^2+^ ion) to macrostate **e** (fully open TM4-TM6 and TM4-TM2 interfaces, destabilized Ca^2+^ ion) are identified in the table insert and represented by the arrows overlaid on the 2D space of the first two tIC vectors (see Methods and also Figure S5 for macrostate assignment). The thickness of the arrows indicates the relative magnitude of the flux of the pathway. The relative contributions of the top pathways to the total flux values are given in the table.

### The open groove conformations in mTMEM16F are stabilized by the allosteric coupling between the distal Ca^2+^ binding site and the groove region

As our results show that the opening of the groove in mTMEM16F is preceded by the restructuring of the distal Ca^2+^ binding site, we hypothesized that the groove opening is allosterically coupled to the dynamics of this site. To quantify such coupling, we first used the TCF formalism we developed ^42,43^ to provide a quantitative description of how particular states, and transitions between them, are favored or opposed by allosteric coupling between reaction coordinates (i.e., CVs). As we had demonstrated that TCFs could be constructed in the context of MSMs, using the first tIC eigenvectors as CVs and the microstate free energies inferred by the MSM ^42^, we used TCF here to quantify the allosteric coupling between the distal Ca^2+^ binding site and the groove opening.

Figure 5A shows the 2D tICA landscape colored according to the TCF values (ΔΔ*A*(*x*, *y*) calculated at each (x, y) grid point of the space (see Methods)). According to Eq. (1), the state (x, y) of the tICA space with ΔΔ*A*(*x*, *y*) < 0 is stabilized by the thermodynamic coupling between tIC1 and tIC2 vectors, while the states with ΔΔ*A*(*x*, *y*) > 0 are destabilized by such coupling ^42^. The TCF map on Figure 5A reveals two pronounced regions on the tICA space stabilized by the coupling between tIC1 and tIC2 (marked by blue rectangles *a* and *b*). The allosterically stabilized states include: 1) the states with fully open TM4-TM2 and TM4-TM6 interfaces (region *a*, the location of microstate 10 from Figure 3), in which the Ca^2+^ ion is displaced from its binding site and coordinating contacts with S854 and I857 residues are lost, leading to a reorganization of the site (see Figure S8); and 2) the state with fully open TM4-TM6 interface but closed TM4-TM2 interface and the Ca^2+^ stably coordinated by all four residues in the distal site (region *b*, the location of microstate 3 in Figure 3). These two regions are separated, in the direction of tIC2, by a region of the space which is destabilized by the allosteric coupling between the two tIC vectors (red rectangle). This state corresponds to the conformations with fully open TM4-TM6 interface, destabilized distal Ca^2+^ binding site, but only partially open TM4-TM2 interface (location of microstate 7 in Figure 3). Finally, two other states can be seen to be destabilized by the thermodynamic coupling between the tIC1 and tIC2 vectors: one corresponds to the conformations with closed TM4-TM6 interface but fully open TM4-TM2 interface and destabilized Ca^2+^ ion in the distal binding site (bottom left region of the tICA space marked by red rectangle); and the other corresponds to the conformations with the TM4-TM6 interface in the beginning stages of destabilization, but closed TM4-TM2 interface and intact distal Ca^2+^ binding site (top left region marked by red rectangle).

**Figure 5:**
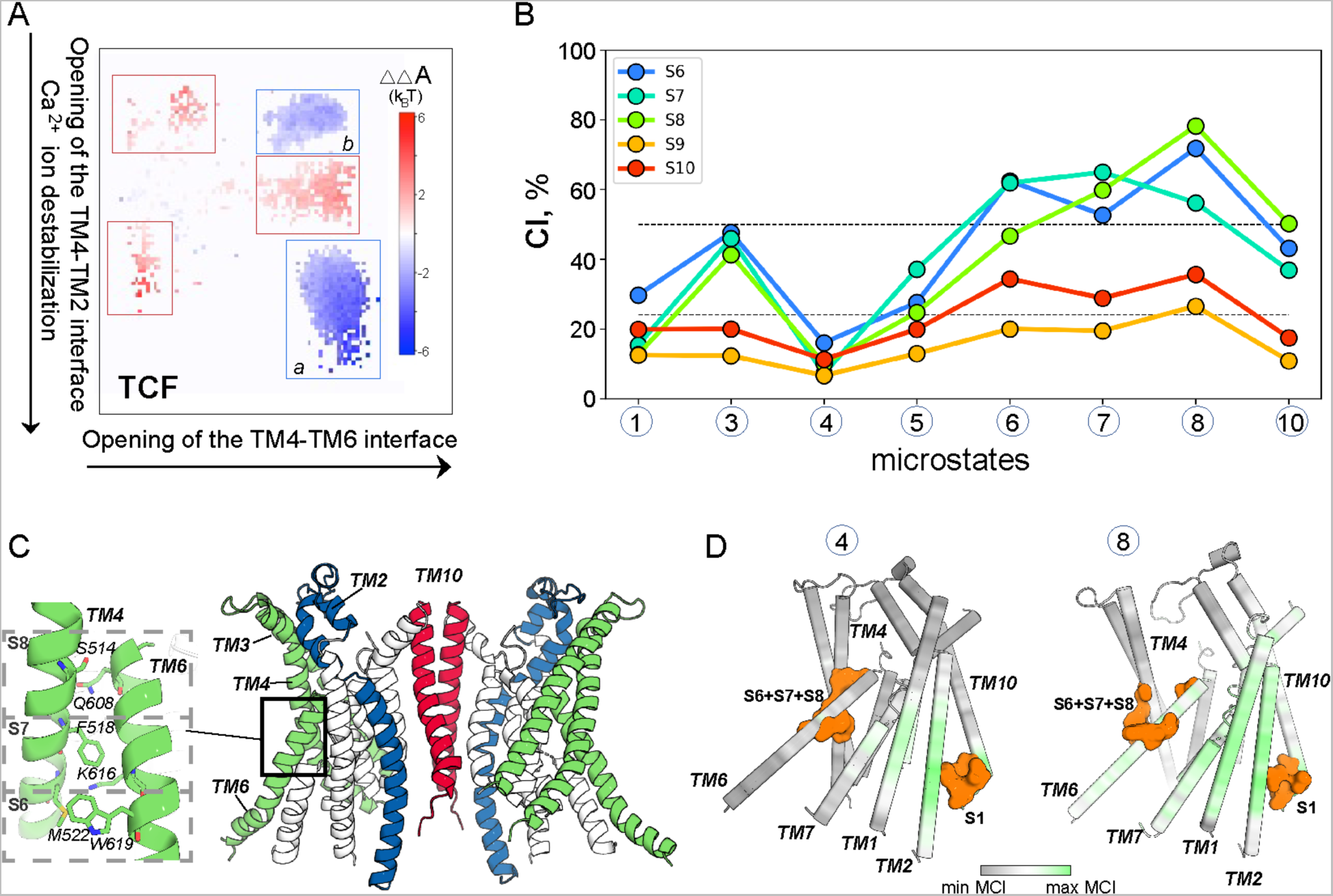
Allosteric coupling between the distal Ca^2+^ binding site and the groove region of mTMEM16F. **(A)** The 2D tICA space from Figure 3 is shown colored according to the TCF. The regions that are stabilized by allosteric coupling between tIC1 and tIC2 vectors (ΔΔ*A*(*x*, *y*) < 0) are highlighted by blue rectangles and labeled as *a* and *b*, and the regions that are destabilized by such coupling (ΔΔ*A*(*x*, *y*) > 0) are marked by red rectangles. (**B**) A plot of the CI values calculated between the *transmitter* S1 site (non-hydrogen atoms of the four Ca^2+^ coordinating residues: D395, S854, I857, D859) and several *receiver* sites at the TM4-TM6 interface (S6 site – non-hydrogen atoms of M522-W619 pair of residues; S7 site – non-hydrogen atoms of F518-K616 pair of residues; S8 site – non-hydrogen atoms of S514-Q608 pair of residues), calculated in the trajectories representing the indicated microstates (see also Figure S9). The two horizontal lines demarcate regions of low, average, and high levels of coordination as obtained from the clustering of the *CI* data using the Fisher-Jenks algorithm (see Methods). (**C**) The mTMEM16F dimer structure highlighting the location and composition of the S6, S7, and S8 sites used in the calculations of *CI* shown in panel B. (**D**) The mTMEM16F monomer structure colored according to the normalized MCI values between the S1 site and the structural locus comprised of the residues from S6, S7, and S8 sites calculated for microstate 4 (left) and 8 (right) trajectories. The residue atoms at S1 and S6+S7+S8 sites are shown as orange surface.

The two main mechanistic inferences from these results are: i) under the conditions of destabilized Ca^2+^ ion in the distal site, the only conformations stabilized by the allosteric coupling between tIC1 and tIC2 are those with fully open groove (i.e., both TM4-TM6 and TM4-TM2 interfaces open); and ii) the fully open TM4-TM6 interface can be allosterically stabilized under two conditions – the fully open TM4-TM2 interface and the reorganized distal Ca^2+^ binding site, or closed TM4-TM2 interface and intact distal Ca^2+^ binding site. Indeed, allosteric coupling between tIC1 and tIC2 makes it unfavorable for the fully open TM4-TM6 interface to exist under conditions of destabilized Ca^2+^ ion if the TM4-TM2 interface is only partially open.

### During the groove opening process the dynamics of the distal Ca^2+^ binding site and the groove region are correlated

To reveal the specific residue groups (structural motifs) involved in the pathway of allosteric coupling between the distal Ca^2+^ binding site and the TM4-TM2 and TM4-TM6 interfaces, we used NbIT to calculate the coordination information (*CI*) between various sites (see Methods). The S1 *transmitter* site combined the residues coordinating the distal Ca^2+^ ion, while *receiver* sites in the TM4-TM2 (S2-S5 sites, Figure S9) and TM4-TM6 (S6-S10 sites, Figure 5C) regions contain residue pairs forming interfacial interactions in the closed groove conformation but are disrupted during the conformational change (see Figure S10 and Methods). The *CI* calculations used the trajectories representing the MSM metastable states identified to lie on the path from fully closed to opened conformations of mTMEM16F groove (microstates 1, 4, 5, 6, 7, 8, and 10 in Figure 3), as well as for the trajectory representing microstate 3 which was identified by the TCF analysis as one of the two regions of the tICA space stabilized by the allosteric coupling between tIC1 and tIC2 vectors (Figure 5A).

The *CI* values between the S1 site and the sites along TM4-TM6 interface (S6-S10) are presented in Figure 5B, while the *CI* values between the S1 site and the sites along TM4-TM2 interface (S2-S5) are shown in Figure S9A. As can be seen from Figure 5B, allosteric coordination between the S1 site and the individual sites along the TM4-TM6 interface (S6-S8) is relatively low for the states with closed or partially open TM4-TM6 region (microstates 1, 4, and 5; see also Figure S6), but it becomes high in the metastable states with fully open TM4-TM6 interface (microstates 3, 6, 7, 8, and 10; Figure S6). Especially notable are the high *CI* values in microstate 8 which represents a transition state along the groove opening pathway in which the TM4-TM6 interface is already open, the distal Ca^2+^ ion is fully destabilized, but the TM4-TM2 interface is still not fully open (Figure S6). Interestingly, *CI* values between the S1 site and sites S9 and S10 that combine the residues from S6–S8 sites located either on TM4 or on TM6 (Figure 5C) are markedly lower in all the microstates (Figure 5B). This suggests that the allosteric communication between the distal Ca^2+^ binding site and the TM4-TM6 region is not due to simply translational motion of TM4 and TM6 helices during the groove opening, but rather to changes in the sidechain fluctuations of the residues at the TM4-TM6 interface as a result of the rupturing of TM4-TM6 interactions during the groove opening process.

The analysis of *CI* between the S1 site and the S2-S4 sites along the TM4-TM2 interface (Figure S9) reveals similar trends. Thus, while the coordination between these sites is overall weaker than that found for the S1 site and the sites along TM4-TM6 interface, we find that the *CI* values in Figure S9 reach their peak again for the protein conformations representing microstate 8, highlighting the important role of this transition state for allosteric communication between the distal Ca^2+^ binding site and the groove region. Furthermore, *CI* between the S1 site and a site at the TM4-TM2 interface containing residues only from TM4 helix (S5 site, see Figure S9), showed relatively weak coordination in all the microstates, suggesting that the allosteric communication between the distal Ca^2+^ binding site and the TM4-TM2 region is related to rupturing of the interactions along the TM4-TM2 interface during the groove opening process.

To identify channels of allosteric communication between the distal Ca^2+^ binding site and the groove region, we quantified the *MCI* between the S1 site and the combination of residues in S6, S7, and S8 sites along the TM4-TM6 interface (see Methods). As shown in Figure 5D (right panel), *MCI* values calculated for the microstate 8 trajectory are high not only in the vicinity of the transmitter and receiver sites but also at other structural loci, most notably throughout TM1, TM2, TM6, and TM7 helices, suggesting that the fluctuations throughout the protein are strongly coupled during the transition towards the groove opening. In contrast, calculated from the trajectory of microstate 4 (in which the *CI* shows the Ca^2+^ binding site and the TM4-TM6 region to be weakly coupled to each other) the calculated *MCI* values are markedly lower (Figure 5D, left panel). Together, the results from this NbIT analysis quantify the allosteric coordination of the groove region dynamics to the dynamic changes in the distal Ca^2+^ binding site through specific residue sets, showing the strength of this coordination to reach its peak in the metastable states on the tICA space that represent transition states on the groove opening pathway.

### The open groove conformation in mTMEM16F conducts lipids

The open, membrane-exposed conformations sampled by the mTMEM16F groove in the MD simulations are similar to experimentally obtained open groove structures in the other TMEM16 scramblases (e.g., in nhTMEM16, see Figure S7), suggesting that the observed open groove state in mTMEM16F would support lipid scrambling. Quantification of the overall number of lipid phosphate headgroups inside the groove volume showed that, in general, the open groove in mTMEM16F can accommodate as many as 4 lipid headgroups at the same time (in Figure S6, see “Lipid Count” histograms for the microstates located at tIC1 > 5), a number consistent with that reported from MD simulations of nhTMEM16 ^28,29,31,32^. Importantly, we observed as well multiple scrambling events through the mTMEM16F open groove structure. Figure 6 details the steps of the lipid scrambling process in four separate MD trajectories (see the figure captions for the criteria used to classify a lipid translocation through the groove as a full scrambling event). In 3/4 cases, the lipid scrambled through the open groove in the EC to IC direction (panels A, C, and D, Figure 6), while in the fourth trajectory, it was scrambled in the opposite, IC to EC direction (panel B, Figure 6). As detailed in Figure S6 by the histograms of distances at the TM4-TM6 and TM4-TM2 interfaces, all 4 of the trajectories projected onto the 2D tICA space in Figure 6 visit the regions characterized large distances that keep these interfaces open.

**Figure 6:**
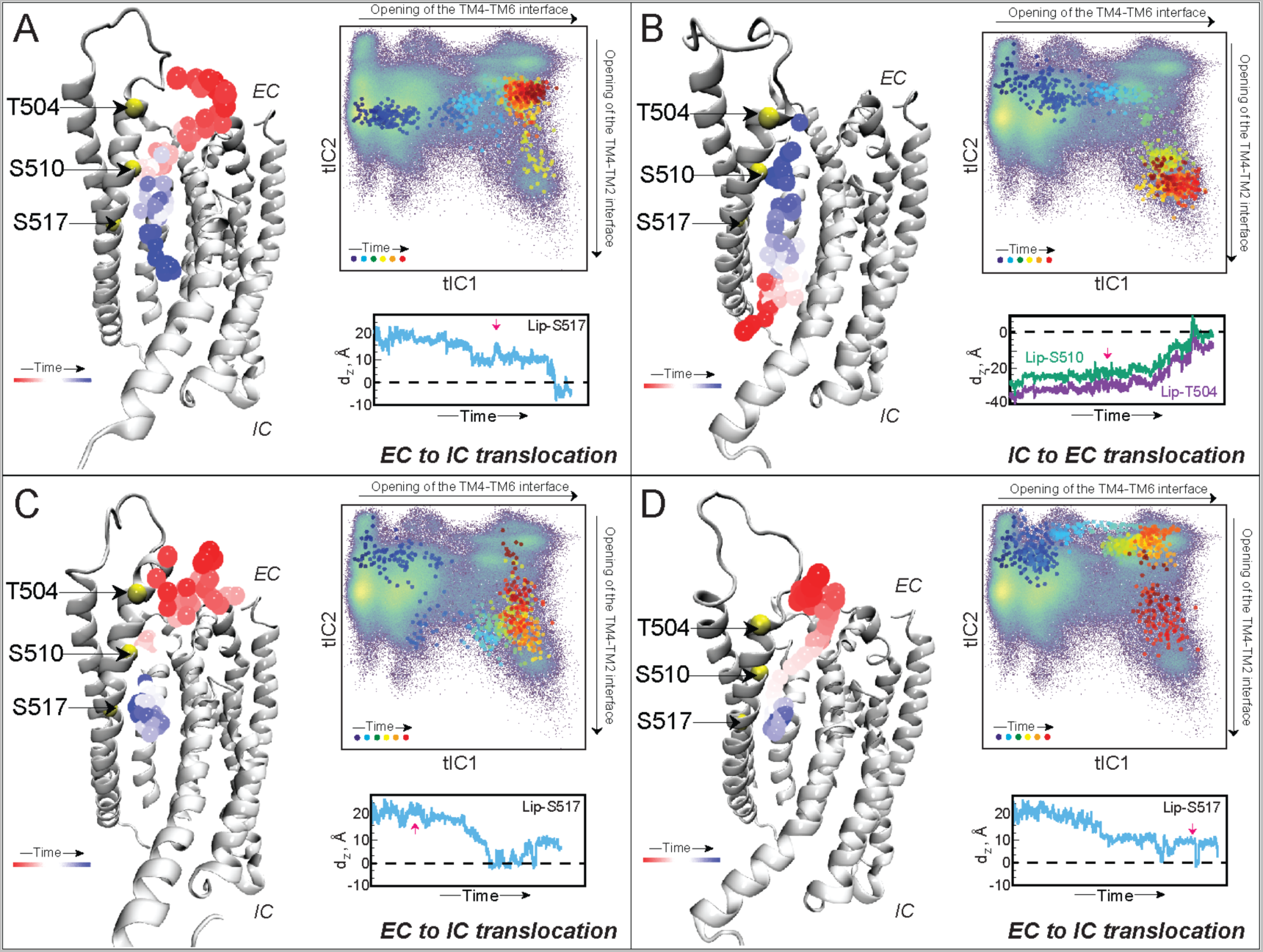
Events of lipid scrambling through the open mTMEM16F groove. The four panels depict lipid scrambling events from four different trajectories. Panels A, C, and D describe lipid translocation from the EC to IC side; panel B describes lipid translocation from the IC to EC side. In each panel, the protein monomer is the structure taken from the final frame of the respective simulation trajectory, shown in cartoon. The trajectory of the scrambled lipid is represented by the phosphorus atom shown as spheres colored according to the timestep (see “Time” color bar). The C_α_ atoms of residues T504, S510, and S517 are shown as yellow spheres. The time evolution of the corresponding MD trajectory projected onto the 2D tICA landscape from Figure 3 is shown as large colored dots with darker colors (blue, cyan) indicating the initial stages of the simulation, lighter colors dots (yellow, green) corresponding to the middle part of the trajectory, and red shades showing the last third of the trajectory. The directions on the tICA space along which TM4-TM6 and TM4-TM2 interfaces open are indicated outside the map, by arrows along the upper border and right border, respectively. The framed time plot in each panel shows the time-evolution of the distance between the phosphorus atom of the scrambled lipid as the Z-distance between the headgroup of the scrambled lipid and the C_α_ atom of residue S517 (d_Z_^S517^, blue trace) in panels A, C, and D. In Panel B, the plot shows the distance of the headgroup and residues T504 and S510 (d_Z_^S504^ and d_Z_^S510^, purple and green traces, respectively). The lipid was considered scrambled in the EC to IC direction if during the trajectory d_Z_^S517^ became < 0, while in the IC to EC direction the lipid was considered scrambled if during the trajectory d_Z_^T504^ became > 0. The red arrows on the Z-distance plots mark the time-points in the trajectory when the TM4-TM2 interface of the groove is fully open (i.e., the system enters microstate 10). The headgroups of POPE and POPG lipid were defined as the center-of-mass of the following group of atoms (in CHARMM36 nomenclature): for POPE – N, C12, C11, P; for POPG – C13, OC3, C12, C11, and P.

As expected from the dynamic properties of the system described above, the initial stages of the IC to EC scrambling process (Figure 6B) are associated with gradual widening of the TM4-TM6 interface and the partial opening of the TM4-TM2 interface; at this point in the trajectory, the translocated lipid is still in the IC part of the groove. Both the TM4-TM6 and the TM4-TM2 interfaces open fully (see red arrow on the d_Z_ plot in Figure 6B) midway through the trajectory, as the system enters microstate 10. Concomitantly, the lipid moves towards the EC vestibule to complete the flipping process (i.e., its headgroup reaches the level of the C_α_ atom of residue T504 located at the very tip of the TM4 helix; see time-evolution of d_Z_ plots). The opening of the TM4-TM2 interface appears to facilitate this last step of the scrambling process. Indeed, in a different trajectory, in which the TM4-TM6 interface fully opened but TM4-TM2 remained closed, a lipid traveling from the IC to EC side performed only a partial flip. This is shown in Figure S11 where the translocated lipid reaches the EC side of the groove and geometrically flips (Figure S10D-E) but remains in the groove at >10Å below the level of the T504 (Figure S11C).

In the three trajectories in which lipid scrambling in EC to IC direction was observed (Figures 5A, C, and D), the opening of the TM4-TM2 interface was seen to facilitate partitioning of a bulk lipid residing in the extracellular leaflet into the groove area to start the translocation process downward towards the IC end. In all three trajectories the TM4-TM2 interface opens partially early in the simulations, as can be inferred from the projections of the trajectories onto the 2D tICA space (locations of dark-colored dots, see also Figure S6). This allows a lipid diffusing near the protein to partition into the EC side of the groove by engaging with positively charged residue K370 on the EC end of TM2 (Figure S12, time-plot “Lip-K370”, and the snapshots at timepoints a, b).

In two of the three trajectories (Figure 6A and 5C), these events are followed by the full opening of both TM4-TM2 and TM4-TM6 interfaces (the systems sample microstate 10). From this point in the trajectory onward the lipid from the EC side starts traveling down the groove to flip (see the red arrow in d_Z_ plots in Figure 6A and 5C). In the third trajectory (Figure 6D), the lipid that entered the EC side of the groove in the early stages of the trajectory moves gradually down the groove and completes the flip after TM4-TM2 interface fully opens (see the red arrow in d_Z_ plot in Figure 6D). In all the trajectories, as the translocated lipid travels through the groove, it is coordinated with another positively charged residue, K616 on TM6 (Figure S10, time-plot “Lip-K616”, and the snapshot at timepoint c).

Together, these results show that lipids can be scrambled through an open groove conformation of mTMEM16F and provide detailed mechanistic descriptions of key steps in the scrambling process.

## DISCUSSION

The results of the computational experiments described here demonstrate that the closed groove structure in mTMEM16F can spontaneously transition into an open, membrane-exposed conformation that supports lipid scrambling. While the properties of open and closed groove conformations in TMEM16F have been previously explored with simulations ^63^, those studies employed structural models built by homology to the known open and closed groove structures of the fungal nhTMEM16 PLS. In contrast, we report here in atomistic detail the dynamic changes that lead from the closed groove structure of mTMEM16F to a scrambling-competent open groove state of this system in MD trajectories totaling ∼400 µs, which allow us to define and quantify the allosteric mechanisms and kinetic model of the process. Thus, we show that the groove opening in mTMEM16F is mechanistically enabled by the allosteric coupling between the two structural motifs that are uniquely present in mTMEM16F but have not been observed together thus far in any available experimentally determined structures of other TMEM16 PLS. One of these motifs is the distal Ca^2+^ ion binding site at the IC end of the dimer interface. The coordination shell of the Ca^2+^ ion at this location includes two acidic residues of which one, E395, resides on TM2. The second structural element involves the extended helix-loop-helix motif which connects the same TM2 helix to TM1 at its other, EC, side. This linker region in mTMEM16F is much longer than the analogous loop region in fungal TMEM16 PLS or in hTMEM16K and reaches over the groove area to contact the EC end of TM4 (Figure S1). This interaction appears to provide additional stability to the closed groove structure in mTMEM16 which is mainly stabilized by contacts between the middle regions of TM4 and TM6 helices.

Remarkably, we find that this situation changes upon the destabilization of the Ca^2+^ ion in its binding site observed in the MD trajectory, which allosterically triggers the disengagement of TM4 from TM2 at their interface. Our analysis showed these two processes to proceed in temporal accord: as the Ca^2+^ ion is displaced from its binding site and moves towards the protein interior, so does the IC part of TM2 as residue D359 maintains the ion coordination. Concomitantly, the EC end of TM2 helix moves away from TM4. These processes are followed by the gradual widening of the TM4-TM6 interface and full disengagement of TM4 from TM2. In this sequence of conformational transitions, the mTMEM16F groove becomes a continuous, membrane-exposed hydrophilic conduit which can be populated, and traversed, by lipid headgroups.

The opening of the TM4-TM2 interface appears to be facilitating the scrambling process from the EC region of the groove in an initial step of groove enlargement that is reminiscent of the mechanistic role of the polar interaction network on the EC ends of TM4 and TM6 helices in nhTMEM16. As we concluded from our MD simulations of the fungal PLS ^29^, the network of polar residue interactions at the EC end of the nhTMEM16 groove, forms a constriction that blocks the passage of lipid headgroups. We showed that dissolution of this interaction network was necessary to trigger full groove opening for lipid translocation through the nhTMEM16 groove ^29^. As mTMEM16F PLS does not contain such interaction network, we propose that a similar mechanistic role can be played here by the interactions between the EC parts of TM4 and TM2. Because the opening of the TM4-TM2 interface is seen to be allosterically connected, through TM2 helix, to the dynamic rearrangements in the distal Ca^2+^ binding site, we suggest that in mTMEM16F the open/close groove equilibrium is regulated by the dynamics of the distal Ca^2+^ ion.

It should be intriguing to establish such a regulatory mechanism in other, as yet structurally uncharacterized TMEM16 scramblases that may contain the two key structural features described above. A possible candidate is TMEM16E, which is highly homologous to TMEM16F (hTMEM16E bears ∼50% sequence identify to mTMEM16F). It is clear, however, that the same type of allosteric communication is unlikely in the fungal TMEM16 PLS, which lack a distal Ca^2+^ ion site, or in hTMEM16K ^35^, in which TM2 does not directly engage in groove stabilization and thus is not expected to be on an allosteric path from the distal Ca^2+^ ion to groove opening.

Because recent studies of the TMEM16A Cl^-^ channel member of the family have shown that Ca^2+^ ion binding at an analogous distal site in this protein was able to facilitate allosterically the channel opening process ^40^, we note that the Ca^2+^ ion coordination shell in TMEM16A consists of three anionic residues in contrast to the coordination shell in mTMEM16F which contains only two anionic sidechains (in TMEM16A, the residue in the position analogous to S854 in mTMEM16F is replaced by Asp). The additional negative charge may entail a different mode of interaction with the protein and Ca^2+^ stability. It will be of interest to compare the allosteric communication between the distal Ca^2+^ binding site and the groove region to the findings presented here.

Thus far, our studies of TMEM16F functional dynamics have not addressed the role of specific lipids, although earlier work had suggested involvement of PIP_2_ lipids in TMEM16F scramblase activity ^34^, and a regulation of scrambling mechanisms of TMEM16 PLS by their lipid environment has been demonstrated ^26,27,39^. The possibility that lipids with different tail lengths and saturation may affect the scrambling by an open-groove state of TMEM16F remains open, as is that of an alternative mechanism of lipid scrambling that envisions lipid translocation with a closed groove^34^. Rather, the open groove scrambling mechanism uncovered in our work may complement the proposed closed groove scrambling mechanism. An interplay between these mechanisms in the context of different lipid environments remains a tantalizing possibility yet to be explored.

## AUTHOR CONTRIBUTION

GK and HW conceived the project. GK designed the study, and with the assistance of XC carried out MD simulations and performed tICA and MSM analyses. EK performed the NbIT analysis. MVL carried out the TCF analysis. All authors contributed to the interpretation of the data and writing the manuscript.

## DECLERATION OF INTERESTS

The authors declare no competing interests.

## ACKNOWLEDGMENTS

We acknowledge stimulating discussions with Prof. Alessio Accardi and his lab members. This work was supported by NIH Grant R01GM106717. H.W. and G.K. gratefully acknowledge support from the 1923 Fund. The authors gratefully acknowledge the enabling role of Chris Carothers, director of the Center for Computational Innovations (CCI) at the Rensselaer Polytechnic Institute (RPI), for the kind help and support of efficient and sustained access to the AiMOS supercomputer at CCI. These resources were generously awarded through the COVID-19 High Performance Computing Consortium. The authors gratefully acknowledge as well the use of resources of the Oak Ridge Leadership Computing Facility, which is a DOE Office of Science User Facility supported under Contract DE-AC05-00OR22725, and of in-house computational resources of the David A. Cofrin Center for Biomedical Information in the Institute for Computational Biomedicine at Weill Cornell Medical College.

## SUPPLEMENTAL FIGURES

**Figure S1:**
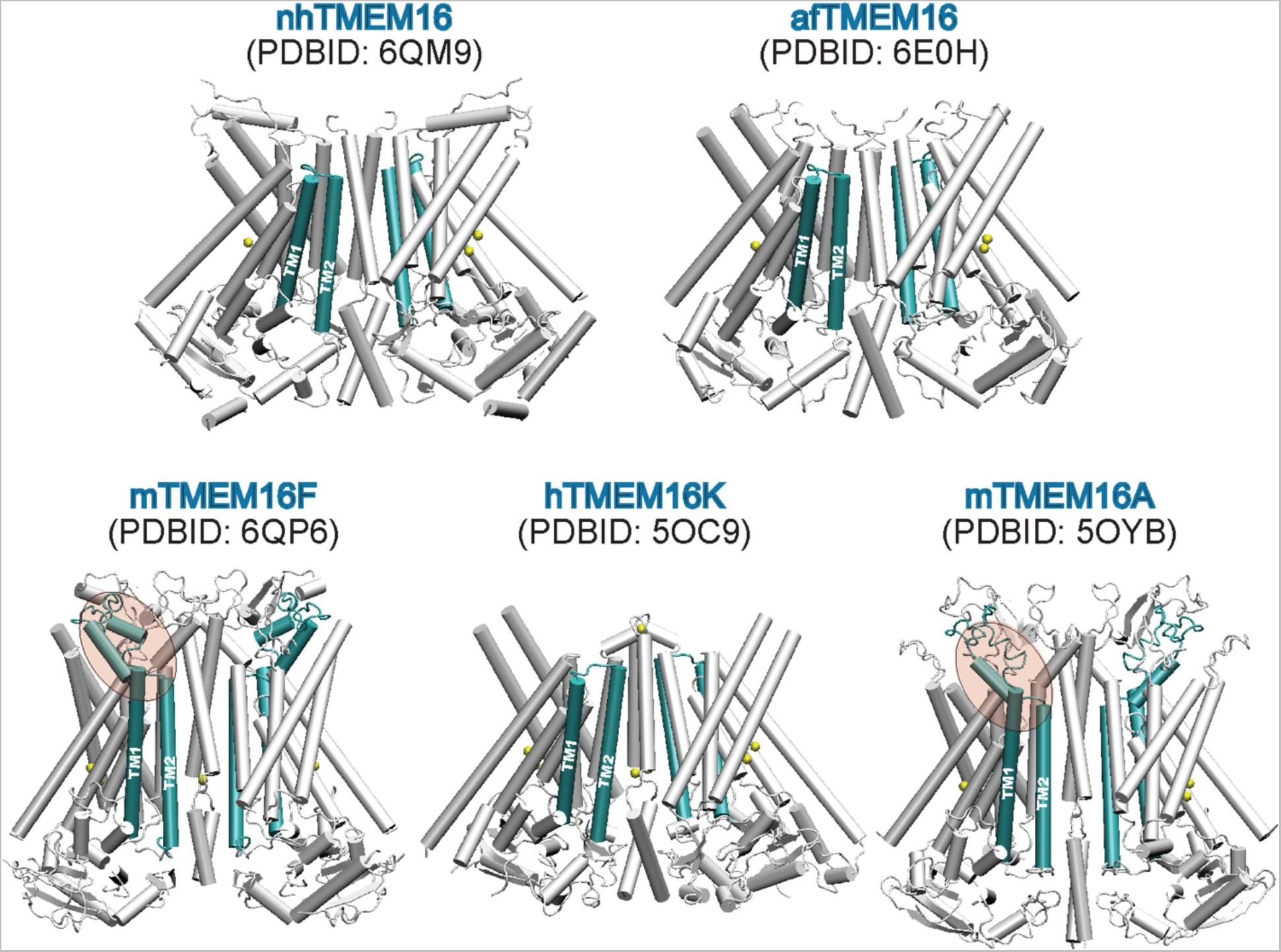
Differences in the characteristics of various structurally determined TMEM16 proteins. Structural comparison of nhTMEM16 (PDBID 6QM9), afTMEM16 (PDBID 6E0H), mTMEM16F (PDBID 6QP6), hTMEM16K (PDBID 5OC9), and mTMEM16A (PDBID 5OYB). TM1-TM2 helices and the connecting loop region are shown in cyan color on each structure. Note that the linker region connecting TMs 1 and 2 in the mTMEM16F and mTMEM16A models reaches towards the groove region (see also Figure 1 in the main text).

**Figure S2:**
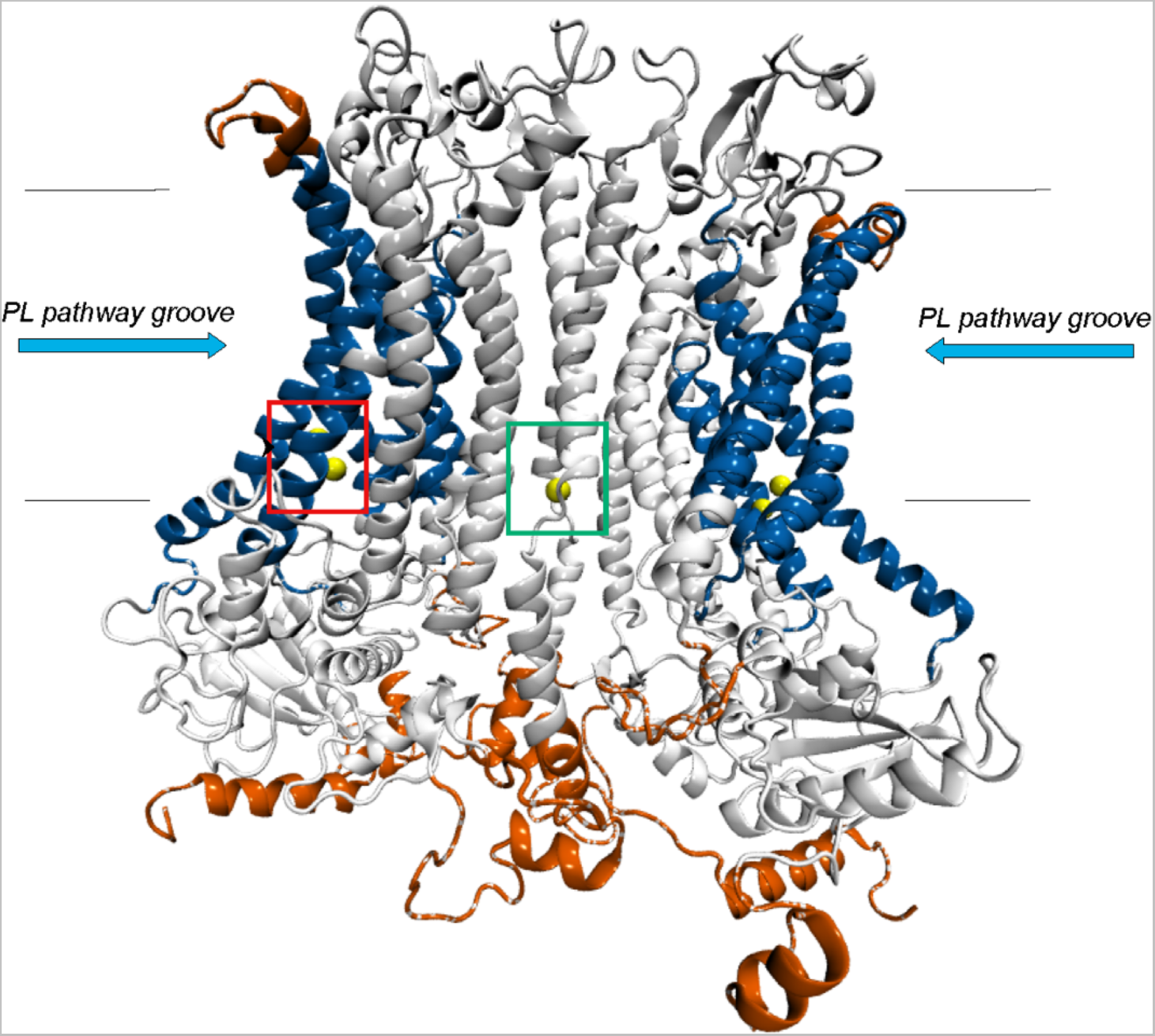
Structural model of the mTMEM16 phospholipid scramblase. The fragments missing from the cryo-EM structure (PDBID 6QP6) were modeled as described and shown in red. The phospholipid (PL) pathway (groove region) is rendered in blue in each protomer. The green box indicates the position of the 3^rd^, distal Ca^2+^ binding site at the dimer interface (all Ca^2+^ ions are shown as yellow spheres). Horizontal black lines demarcate the relative positions of upper and lower leaflets of the membrane (i.e., idealized positions of phosphate groups on the two leaflets).

**Figure S3:**
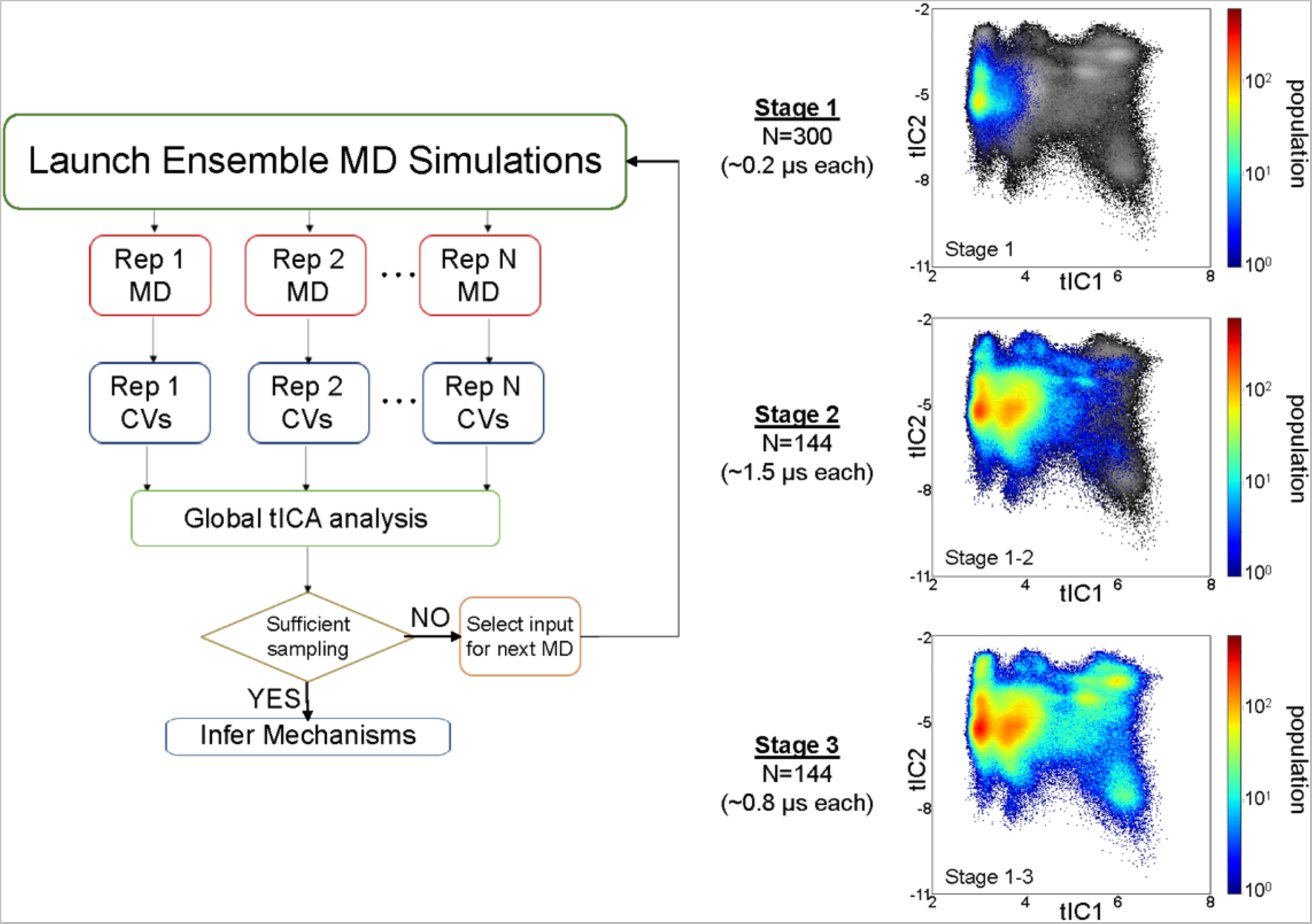
Protocol of the multi-stage adaptive ensemble MD simulation. ***Left panel:*** Flowchart detailing steps of the protocol. Each stage of the adaptive ensemble MD simulations consists of 1) running *N* number of independent replicates of the system; 2) Calculating collective variables (CVs) from each trajectory; and 3) Combining the CVs into a global tICA dimensionality reduction analysis. In each row, Rep 1 to Rep N represent the individual replicas run separately. If the sampling of the phase space is satisfactory, subsequent analyses are performed to infer kinetic and molecular mechanisms. If more sampling is required, a next round of ensemble MD simulations is launched from selected frames from the previous stage (see Methods for more details). Here, this adaptive protocol was carried out in 3 stages. The number of independent replicates for each state and lengths of MD simulation runs for each replicate in each stage are shown to the right of the Flowchart. ***Panels on the right:*** Projections of the frames from the trajectories obtained at each stage onto the final (Stage 3) 2D tICA space spanned by the first 2 tIC vectors – tIC1 and tIC2 (see Methods for details). The populations of the different states on the final tICA space are indicated by the colored portions in each stage, superimposed on a greyscale representation of the Stage 3 tICA space. Together, the three panels show the gradual increase of the tICA space sampling as the adaptive protocol is executed in 3 stages.

**Figure S4:**
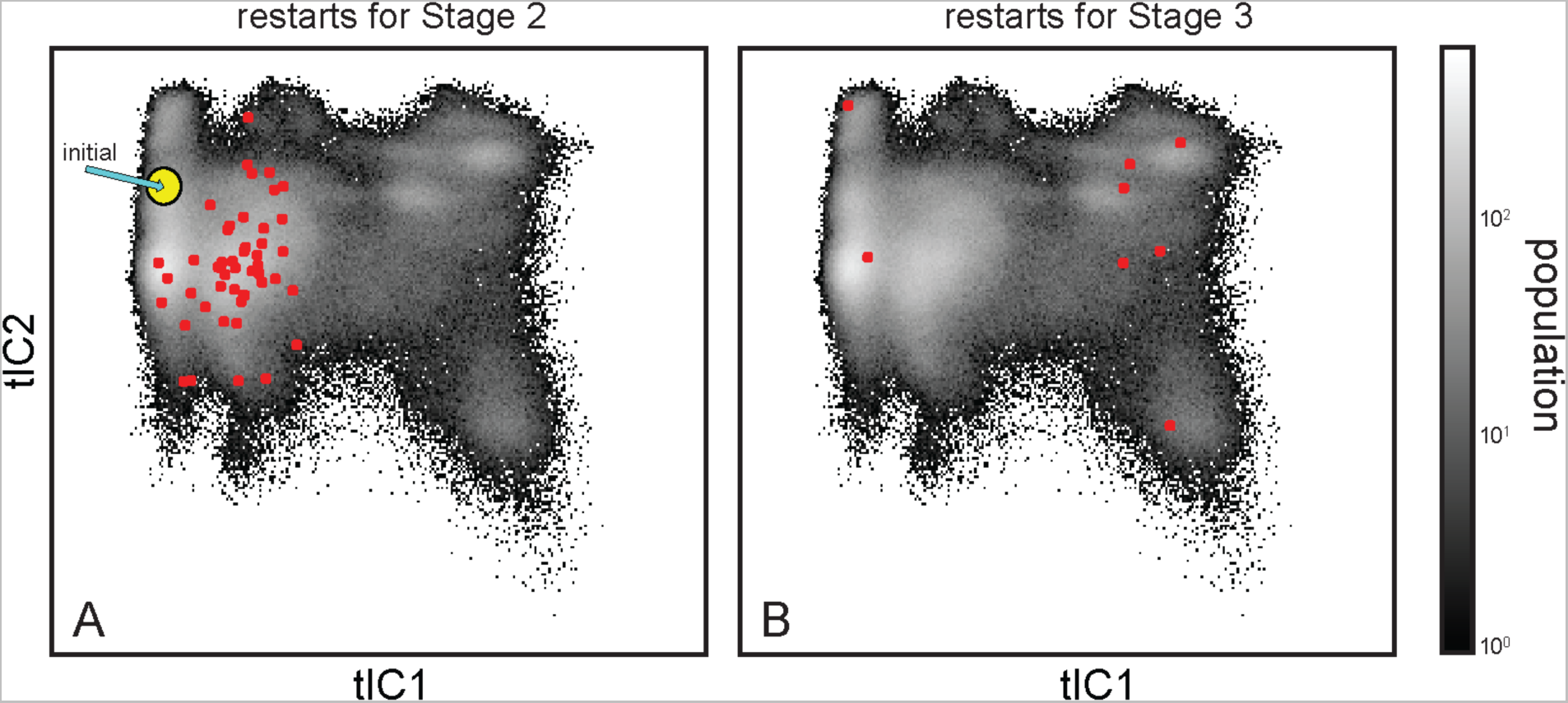
Restart frames of trajectories started in each stage of the adaptive ensemble MD simulation protocol. The restart frames are shown as red circles projected onto the final 2D tICA landscape shown in grayscale, for (**A**) Stage 1 and (**B**) Stage 2 simulations (see also Figure S3). The location on the tICA space of the initial, cryo-EM structure is also indicated in panel A by the yellow circle.

**Figure S5:**
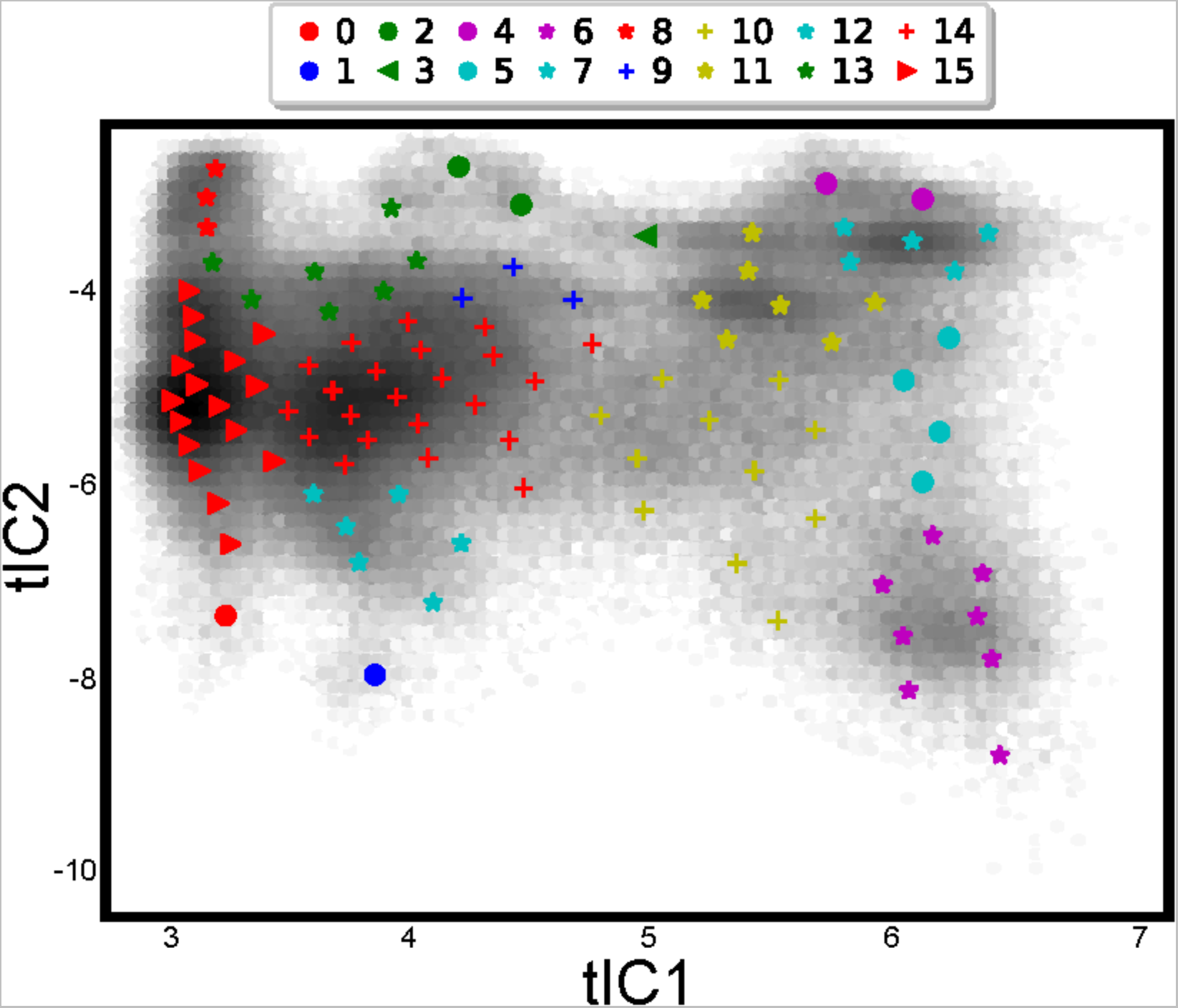
Discretization of the 2D space of the first two tIC vectors into structurally similar microstates and kinetically similar macrostates. The 2D space of tIC1 and tIC2 vectors (in grey) is discretized into 100 microstates. The centroids of these structurally similar microstates are shown as symbols of various colors and sizes, according to their grouping into 16 kinetically similar macrostates (see Methods).

**Figure S6:**
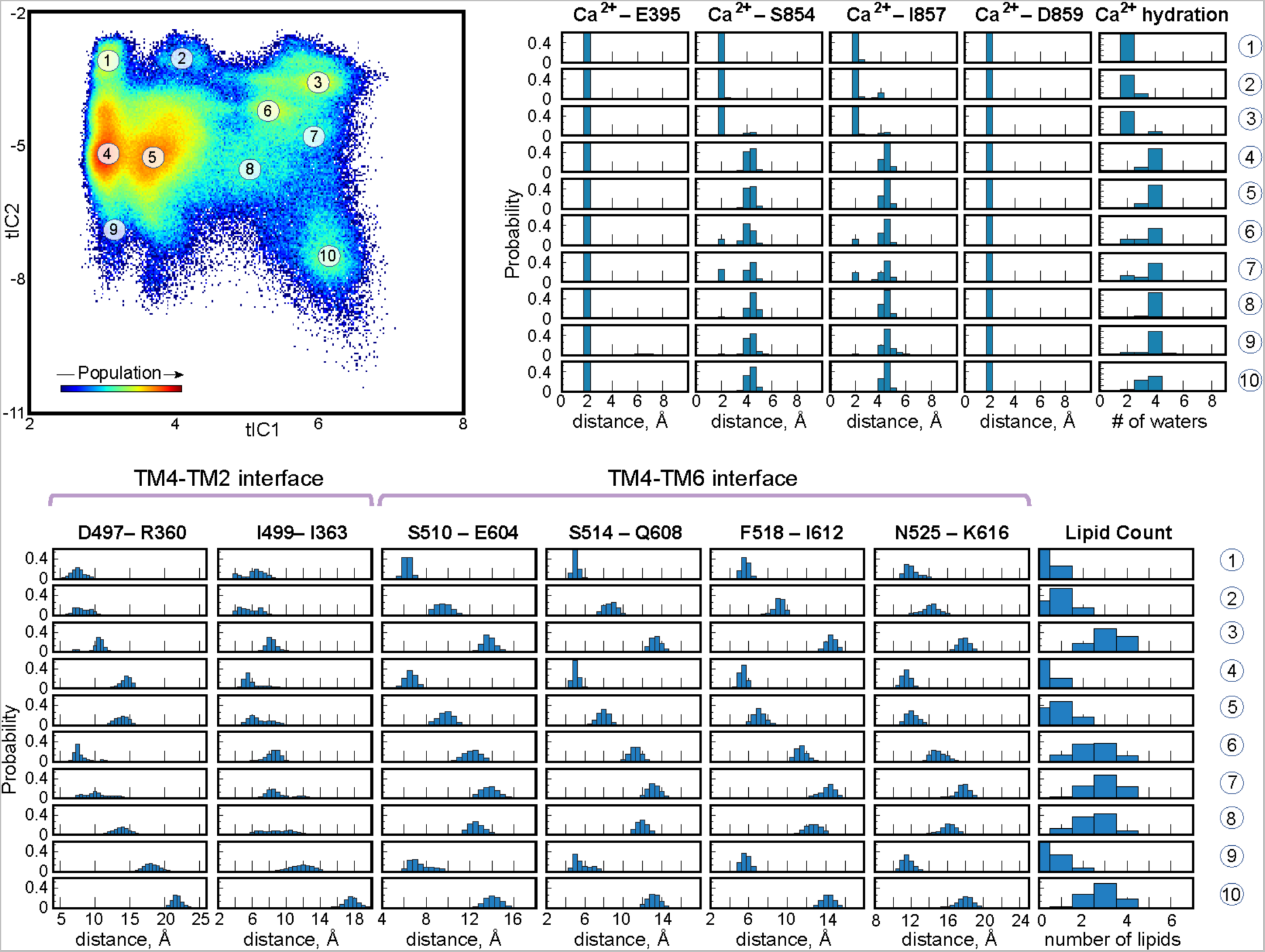
Structural characteristics of various microstates on the tICA space. The positions of selected microstates are indicated on the 2D space of the first two tIC vectors (as in Figure 3). The structural characteristics of the TM4-TM6 interface, TM4-TM2 interface, and the Ca^2+^ ion binding site are quantified in the accompanying histograms for each of the selected microstates. The # of water molecules entry is defined as the number of oxygen atoms at a distance of 3Å from the distal Ca^2+^ ion. The number of lipids entry is defined as the number of lipid phosphorus atoms within 7Å of the indicated residue.

**Figure S7:**
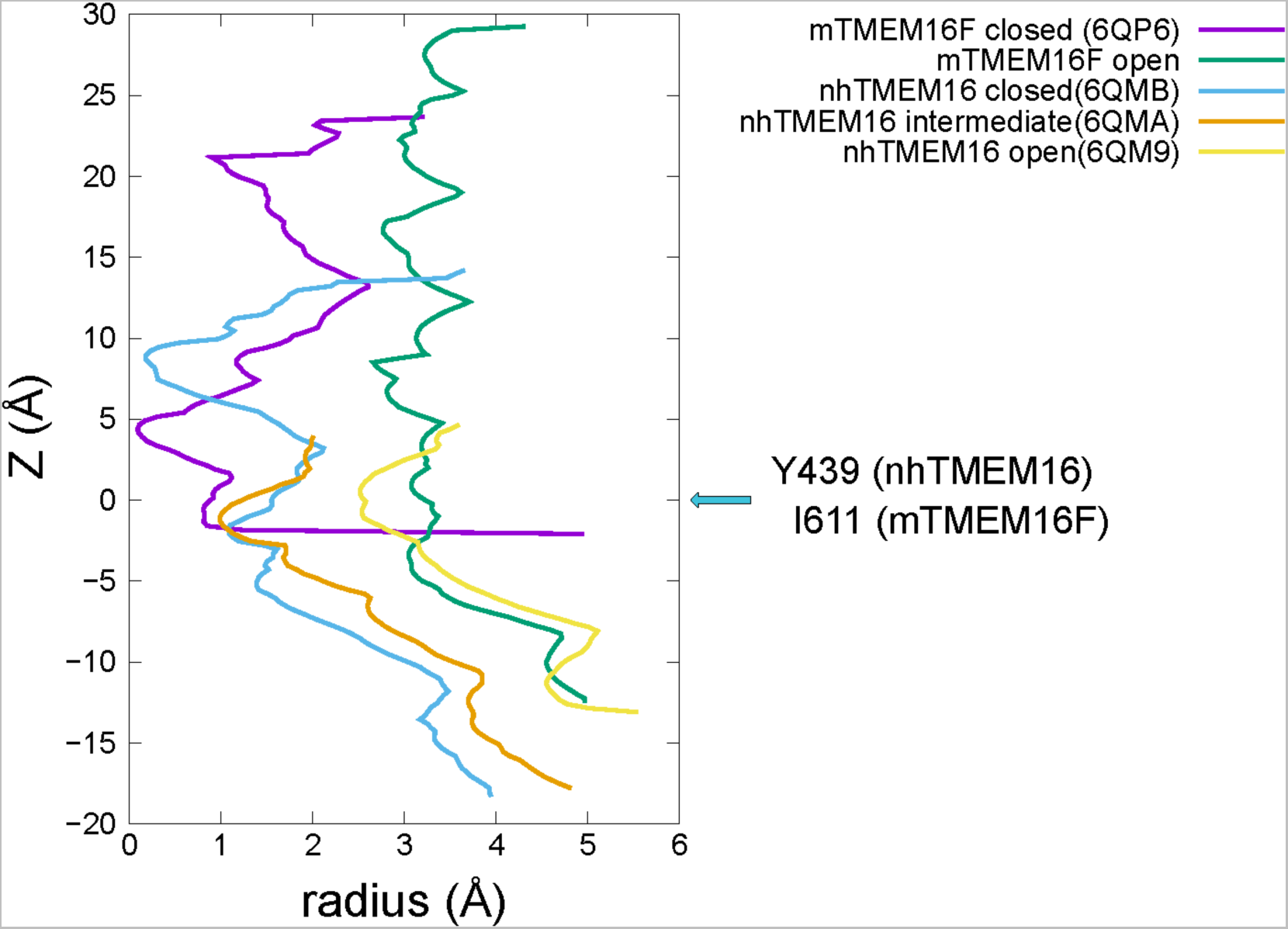
Comparison of pore dimensions in the open groove conformation of mTMEM16F to the open groove structure of nhTMEM16. The radius of the permeation pathway of nhTMEM16 and mTMEM16F was estimated using the program HOLE (http://www.holeprogram.org/) for the various states, as indicated. Purple: Closed groove structure of mTMEM16F (PDBID 6QP6); Green: Open groove structure of mTMEM16F (representative snapshot from microstate vii in Figure 3); Blue: Closed groove structure of nhTMEM16 Open (PDBID 6QMB); Orange: Intermediate groove structure in nhTMEM16 (PDBID 6QMA); Yellow: Open groove structure of nhTMEM16 (PDBID 6QM9). The the cyan-color arrow marks the Z=0Å coordinate and indicates the position of the reference residue in nhTMEM16 (Y439) and mTMEM16F (I611) along the z axis.

**Figure S8:**
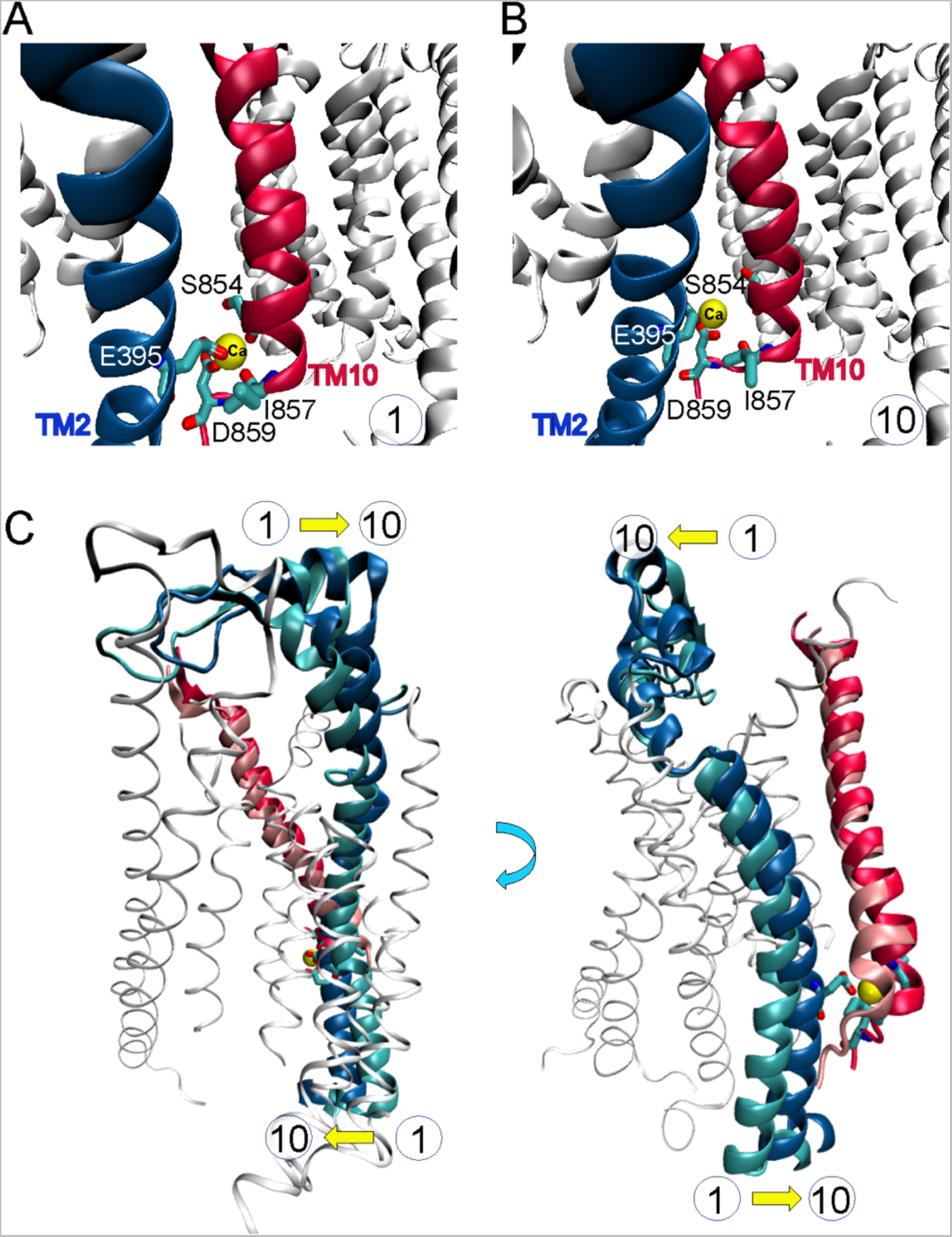
Conformational changes in the distal Ca^2+^ ion binding site at the mTMEM16F dimer interface. **(A)** Ca^2+^ ion (yellow sphere) is stably bound in the binding site, coordinated by the sidechains of E395 and D859 residues, as well as by the backbone carbonyls of S854 and I857. This interaction mode is representative of *microstate 1* in Figure 3. **(B)** Ca^2+^ ion is destabilized in the binding site. It has lost contact with the backbone carbonyls and is now coordinated solely by the sidechains of E395 and D859 residues. This interaction mode is representative of *microstate 10* in Figure 3. TM2 and TM10 helices where the coordinating residues reside are shown in blue and red color, respectively. **(C)** Two views of the mTMEM16F monomer highlighting conformational changes in TM2 and TM10 helices during the transition from *microstate 1 to 10*. The conformations of TM2 in microstates 1 and 10 are shown in cyan and blue color, respectively, whereas conformations of TM10 helix in microstates 1 and 10 are depicted in pink and red, respectively. The conformation of the rest of the protein structure is from microstate 10.

**Figure S9:**
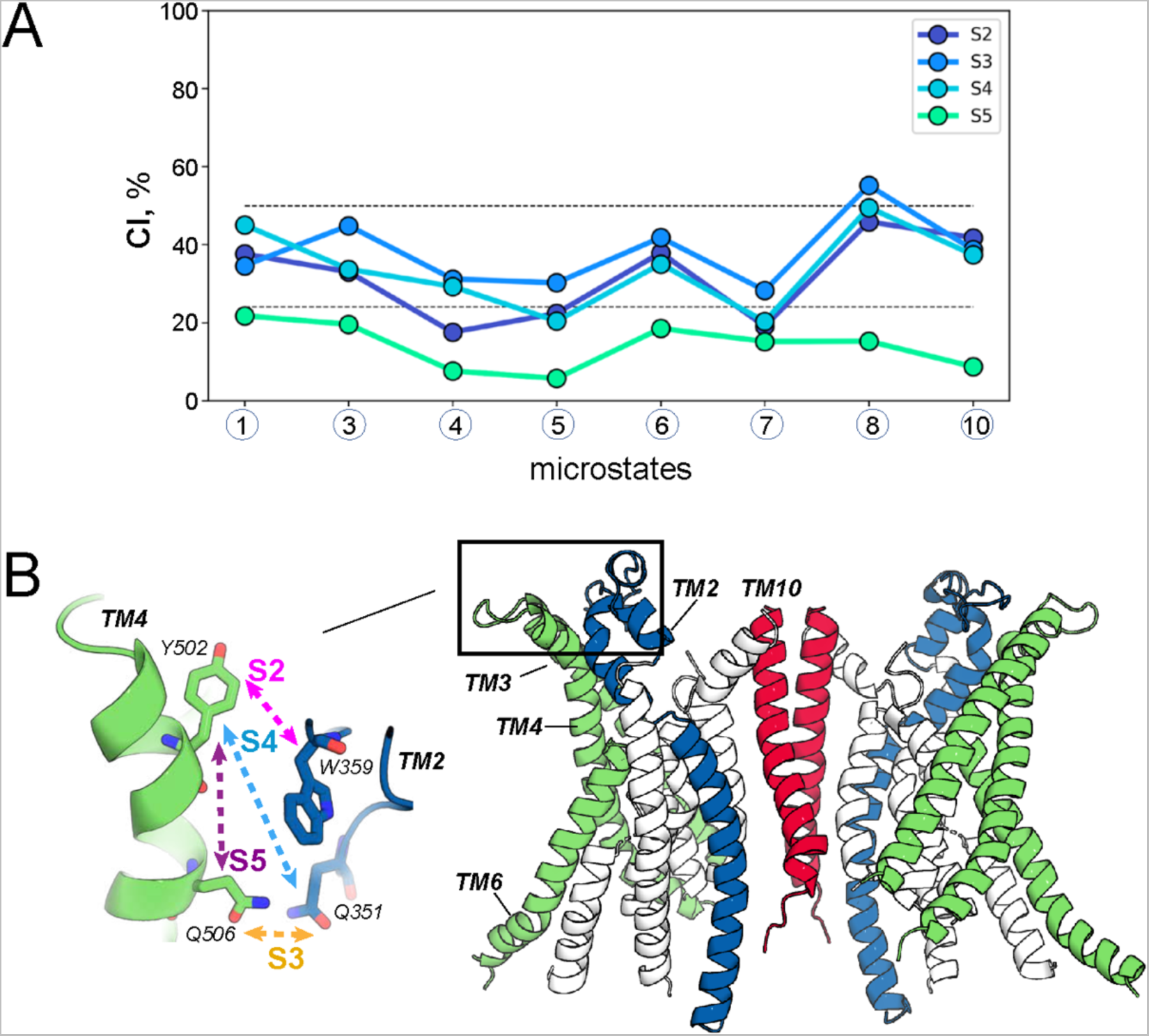
Allosteric coupling between the distal Ca^2+^ binding site and the TM4-TM2 interface of the groove region in mTMEM16F. (**A**) The coordination information (**CI**) between site S1 (non-hydrogen atoms of the four Ca^2+^ coordinating residues: D395, S854, I857, D859) and several sites at the TM4-TM2 interface (S2 site – non-hydrogen atoms of Y502-W359 pair of residues; S3 site – non-hydrogen atoms of Q504-Q351 pair of residues; S4 site – non-hydrogen atoms of Y502-Q351 pair of residues; S5 site – non-hydrogen atoms of Y502-Q506 pair of residues) were calculated from the trajectories representing the indicated microstates (see also Figure 5). The horizontal lines demarcate regions of *low*, *average*, and *high* levels of coordination as obtained from the clustering of the *CI* data using the Fisher-Jenks algorithm (see Methods). (**B**) The mTMEM16F dimer structure highlighting the location and composition of the S2, S3, S4, and S5 sites used in the calculations of *CI* shown in panel A.

**Figure S10:**
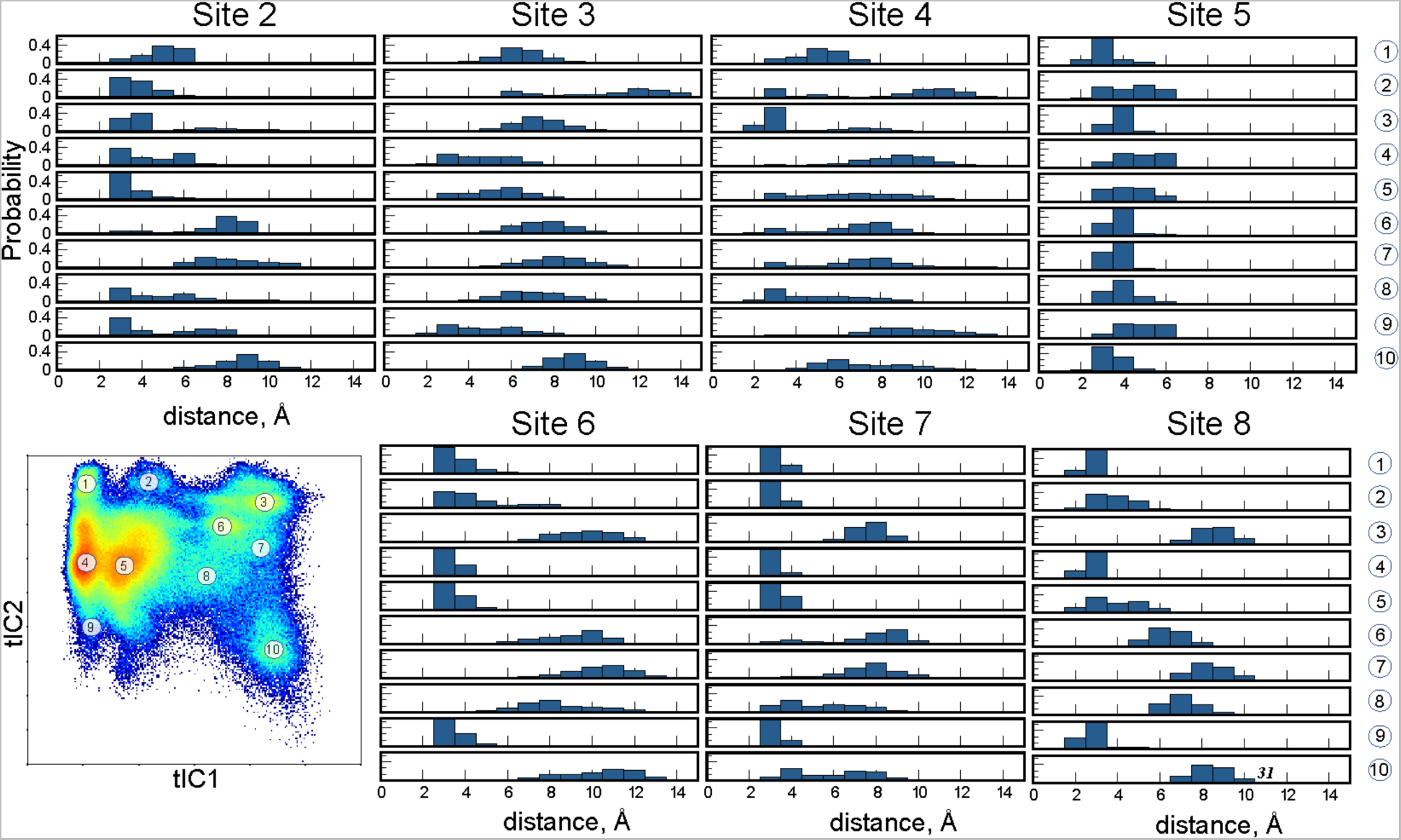
Structural characteristics of various NbIT sites. Pair-wise minimum distances of the residues at NbIT (*receiver*) Sites 2-8 in selected microstates of the 2D tICA space. S2 site – W359/Y502, S3 site – Q506/Q351; S4 site – Q351/Y502; S5 site – Y502/Q506; S6 site – M522/W619; S7 site – F518/K616; S8 site – S514/ Q608; S9 site – S514/F518/M522; S10 site – Q608/K616/W619.

**Figure S11:**
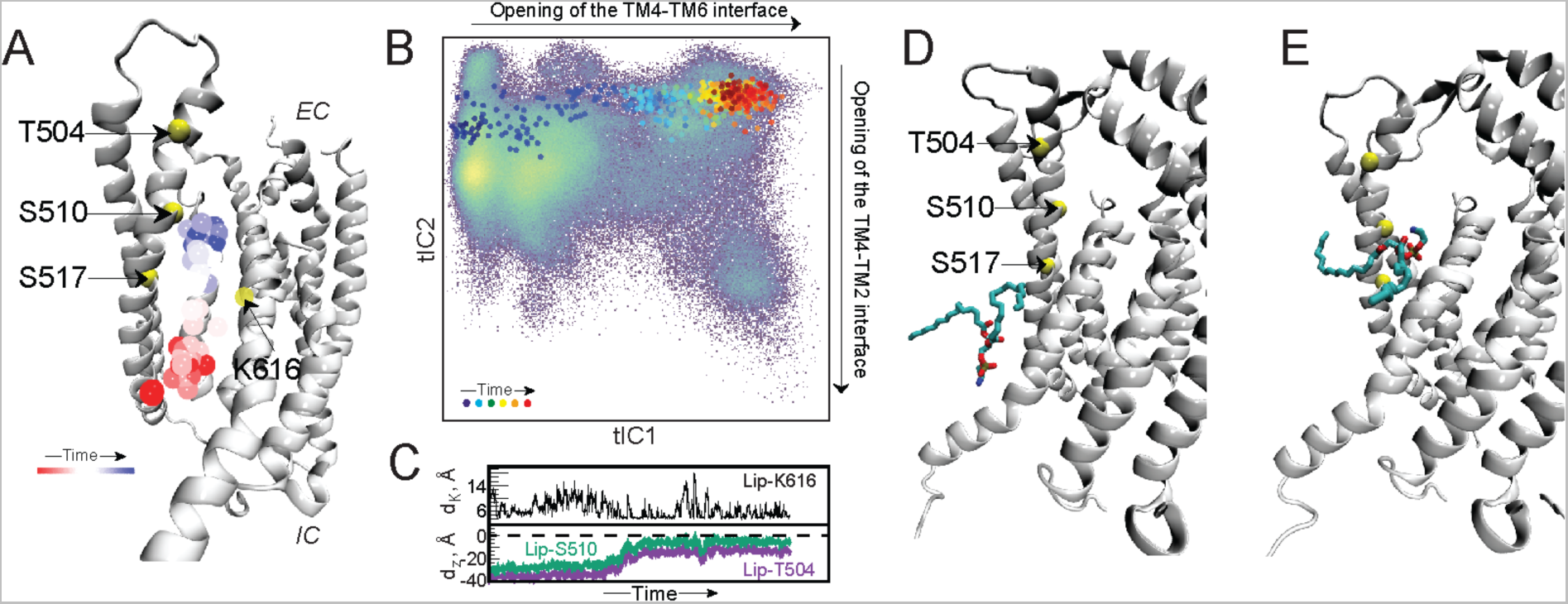
Incomplete lipid scrambling event in the MD simulations. (**A**) The protein monomer is shown (in cartoon, the structure taken from the final frame of the respective simulation trajectory) and the trajectory of the phosphorus atom of the translocated lipid (from the IC to EC side) is shown as spheres colored according to the timestep (see “Time” color bar). The C_α_ atoms of residues T504, S510, and S517 are shown as yellow spheres. (**B**) The time evolution of the corresponding MD trajectory projected onto the 2D tICA landscape from Figure 3 is shown as large colored dots. The darker colors (blue, cyan) indicate the initial stages of the simulation, lighter colors dots (yellow, green) indicate the middle part of the trajectory, and red shades show the last third of the trajectory. The directions on the tICA space along which TM4-TM6 and TM4-TM2 interfaces open are indicated. (**C**) The time-evolution of the distance between the phosphorus atom of the scrambled lipid and the amine nitrogen atom of residue K616 sidechain is shown (d_K_, black traces), and the time-evolution of the Z-distance between the headgroup of the translocated lipid and the C_α_ atom of residues T504 and S510 (d_Z_^S504^ and d_Z_^S510^) is shown by the purple and green traces, respectively. (**D-E**) Snapshots from the first (**D**) and the last (**E**) frames of the trajectory showing the positioning of the translocated lipid. The color code for the protein is the same as in panel **A**. Note that, while the lipid is geometrically flipped during the simulation, it is still bound in the groove as shown by the d_Z_^S517^ distance remaining < 0 (see panel **B**). The headgroups of POPE and POPG lipids were defined as the center-of-mass of the following group of atoms (in CHARMM36 nomenclature): POPE (N, C12, C11, P); POPG (C13, OC3, C12, C11, P).

**Figure S12:**
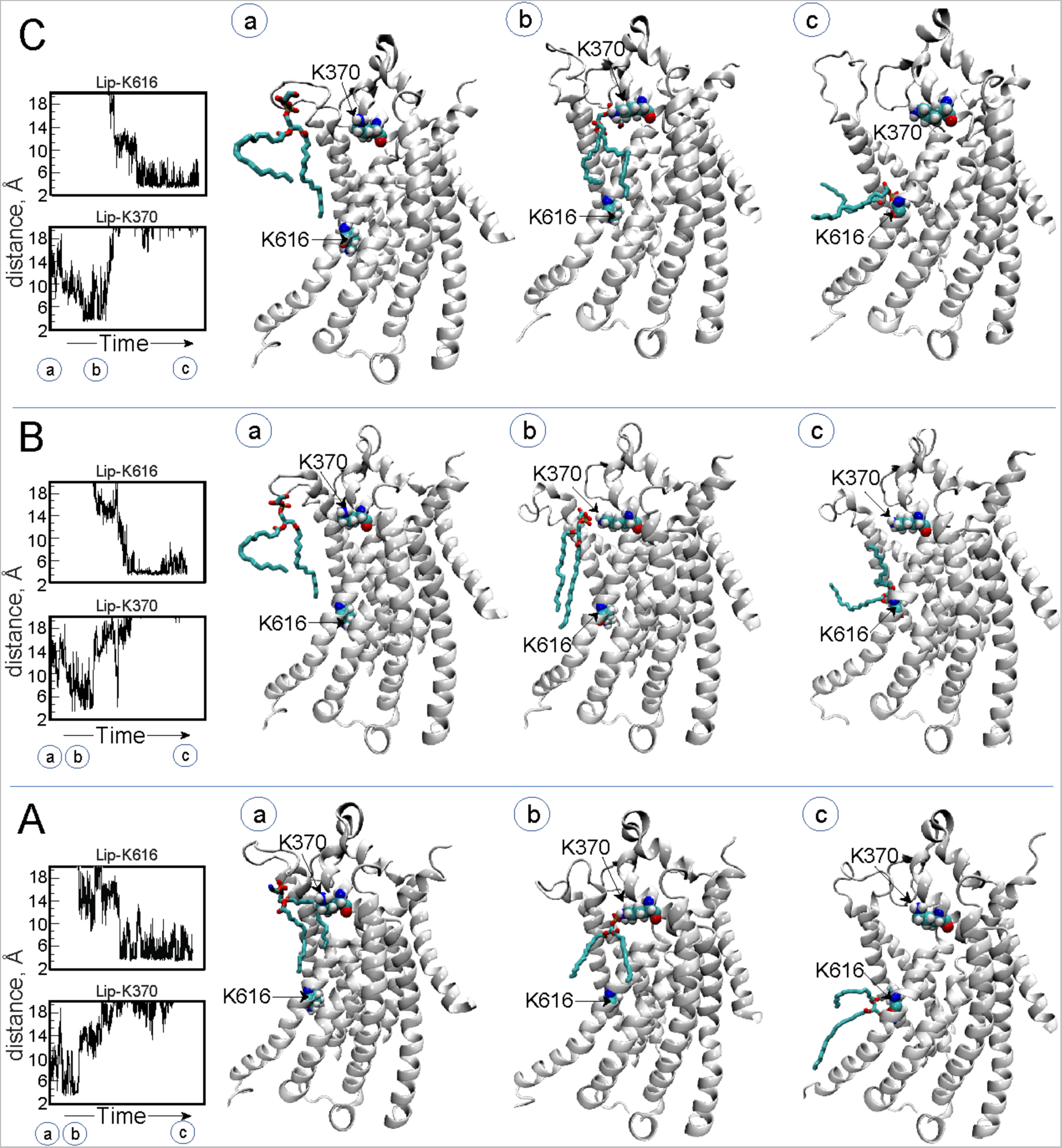
The scrambled lipid is coordinated by basic residues in the groove area. Panels **A, B,** and **C** show coordination by the positively charged residues of the lipid translocated from the EC to IC side of the groove in the three trajectories described in Figure 5 (panels A, B, and D, respectively). In each panel, the left graphs show the time-evolution of the distance between the phosphorus atom of the scrambled lipid and the nitrogen of the amine group of residues K616 and K370. The snapshots show the configuration of the system at three specified time points (marked as **a, b,** and **c**) along the respective trajectory. In these snapshots, the mTMEM16F monomer is drawn in white cartoon, the scrambled lipid is in licorice and two basic residues in the groove area, K616 and K370 are shown in space-fill representation.

